# Loss of RREB1 in pancreatic beta cells reduces cellular insulin content and affects endocrine cell gene expression

**DOI:** 10.1101/2022.06.04.494826

**Authors:** Katia K Mattis, Nicole AJ Krentz, Christoph Metzendorf, Fernando Abaitua, Aliya F Spigelman, Han Sun, Antje K Rottner, Austin Bautista, Eugenia Mazzaferro, Marta Perez-Alcantara, Jocelyn E Manning Fox, Jason M Torres, Agata Weslowska-Andersen, Grace Z Yu, Anubha Mahajan, Anders Larsson, Patrick E MacDonald, Benjamin Davies, Marcel den Hoed, Anna L Gloyn

## Abstract

**Aims/hypothesis:** Genome-wide studies have uncovered multiple independent signals at the *RREB1* locus associated with altered type 2 diabetes risk and related glycemic traits. However, little is known about the function of the zinc finger transcription factor RREB1 in glucose homeostasis or how changes in its expression and/or function influence diabetes risk.

**Methods:** A zebrafish model lacking *rreb1a* and *rreb1b* was used to study the effect of RREB1 loss *in vivo*. Using transcriptomic and cellular phenotyping of a human beta cell model (EndoC-βH1) and human induced pluripotent stem cell (hiPSC)-derived beta-like cells, we investigated how loss of RREB1 expression and activity affects pancreatic endocrine cell development and function. *Ex vivo* measurements of human islet function were performed in donor islets from carriers of *RREB1* T2D-risk alleles.

**Results:** CRISPR-Cas9-mediated loss of *rreb1a* and *rreb1b* function in zebrafish supports an *in vivo* role for the transcription factor in beta cell mass, beta cell insulin expression, and glucose levels. Loss of RREB1 reduced insulin gene expression and cellular insulin content in EndoC-βH1 cells, and impaired insulin secretion under prolonged stimulation. Transcriptomic analysis of *RREB1* knockdown and knockout EndoC-βH1 cells supports RREB1 as a novel regulator of genes involved in insulin secretion. *In vitro* differentiation of *RREB1*^KO/KO^ hiPSCs revealed a dysregulation of pro-endocrine cell genes, including *RFX* family members, suggesting that RREB1 also regulates genes involved in endocrine cell development. Human donor islets from carriers of T2D-risk alleles in *RREB1* have altered glucose-stimulated insulin secretion *ex vivo*, consistent with RREB1 regulating islet cell function.

**Conclusions/interpretation:** Together, our results indicate that RREB1 regulates beta cell function by transcriptionally regulating the expression of genes involved in beta cell development and function.

**Research in context:** *What is already known about this subject?:*

- Human genetic variation in *RREB1* is associated with altered diabetes risk, variation in glycemic, and anthropometric traits
- RREB1 is a transcription factor that binds to Ras-responsive elements and is expressed in multiple diabetes relevant tissues, including pancreatic islets

*What is the key question?:*

- How does altered expression or function of RREB1 influence diabetes risk?

*What are the new findings?:*

- Knockdown and knockout of *RREB1* in mature human EndoC-βH1 cells reduces expression of insulin transcript and cellular content, as well as insulin secretion under prolonged stress
- Carriers of the T2D-risk *RREB1* coding allele trend towards reduced insulin content, but have improved glucose-stimulated insulin secretion
- A loss-of-function zebrafish model suggests that RREB1 is required for insulin expression

*How might this impact on clinical practice in the foreseeable future?:*

- RREB1 controls beta cell function and whole-body glucose homeostasis by transcriptionally regulating the development and function of pancreatic beta cells

## Introduction

Genome-wide association studies have discovered multiple independent signals at the *RREB1/SSR1* locus that are associated with altered type 2 diabetes (T2D)-risk and various metabolic and anthropometric traits, including fasting glucose levels and height [1–4]. Genetic fine-mapping identified the coding variant rs9379084 (p.Asp1171Asn) as causal (92% posterior probability for T2D), strongly supporting *RREB1* as the effector transcript at this locus [5]. Carriers of the minor allele encoding p.Asn1171-*RREB1* - predicted to have a detrimental effect on RREB1 protein function (CADD-score 28.2) - have a lower risk of developing T2D and lower fasting glucose levels, on average [2]. The shared association between T2D risk and quantitative measures of islet function supports the islet as a key tissue mediating disease [6–8], and suggests a potential role for RREB1 in beta cell development and/or function.

Although RREB1 has been studied in several different cellular contexts, there have been no investigations on its role in the pancreatic beta cell [9–11]. *RREB1* encodes a zinc finger transcription factor that is expressed in several T2D-relevant tissues, including pancreatic islets, adipose tissue, liver and skeletal muscle [12–14]. Several lines of evidence support a potential developmental role for RREB1: 1) Homozygous deletion of *Rreb1* in mice is embryonic lethal [10]; 2) *RREB1* transcript [15] and protein [16] is detected during *in vitro* endocrine cell differentiation of human induced pluripotent stem cells (hiPSC); and 3) RREB1 is a downstream target of the MAPK/ERK, a signalling pathway that is important for early human beta cell differentiation [17]. However, whether genetic variation in *RREB1* influences diabetes risk due to altered endocrine cell development and/or function is unknown.

Characterisation of an *in vivo* zebrafish model deficient in orthologues *rreb1a* and *rreb1b* revealed an increased beta cell mass with reductions in average beta cell insulin expression, glucose levels, and length. To explore the role of RREB1 in endocrine cell development and beta cell function, we generated both isogenic *RREB1* knockout and wildtype hiPSC lines, as well as non-clonal *RREB1* knockout EndoC-βH1 mature beta cells. Comprehensive transcriptomic and cellular phenotyping of our human beta cell models demonstrated a role for RREB1 in both mature beta cell function and endocrine cell development, and the transcriptional regulation of *RFX* family members. Consistent with the cellular models, there were significant differences in glucose-stimulated insulin secretion in human donor islets from carriers of *RREB1* T2D-risk alleles compared to non-carriers. Together, we have identified a transcriptional role for RREB1 that is consistent with genetic variation in *RREB1* influencing diabetes risk through effects on the pancreatic islet cell development.

## Methods

### Zebrafish studies

#### Husbandry and transgenic lines

Adult zebrafish (*Danio rerio*) were housed in systems with recirculating, filtered water of 28.5°C (Aquaneering Inc, USA) on a 14/10 h light/dark cycle. Through crossing we generated AB fish with transgenically expressed, fluorescently labelled pancreatic beta cell nuclei (*tg(−1.2ins:H2B-mCherry)*) [18] and hepatocytes (*tg(fabp10a:EGFP)*) [19, 20]. AB fish for outcrossing and line maintenance were obtained from the European Zebrafish Resource Center (EZRC); beta cell and liver reporter lines were kindly provided by Vanderbilt University (*tg(−1.2ins:H2B-mCherry)*) and the EZRC (*tg(fabp10a:EGFP)*). All zebrafish handling and experiments were carried out in agreement with Swedish animal welfare laws and were approved by the Uppsala University Ethical Committee for Animal Research (Dnrs C14/16 and 5.8.18-13680/2020).

#### Sequence analysis

Human *RREB1* amino acid sequences and those of the zebrafish orthologues (*rreb1a* and *rreb1b*, **ESM Table 1**) were downloaded from uniprot (www.uniprot.org). Amino acid sequences across species were aligned using MUSCLE multiple sequence alignment and a phylogenic tree was constructed using the Neighbor-Joining method with bootstrapping (2000 replicates) within the MEGA11 software [21].

#### CRISPR/Cas9 in zebrafish embryos

*Danio rerio* genome version GRCz11, ENSEMBL.org, was used for planning all CRISPR/Cas9-related work. Suitable guide RNAs in the coding regions of the *rreb1a* (ENSDARG00000063701) and *rreb1b* (ENSDARG00000042652) genes were identified using CRISPOR (http://crispor.tefor.net; [22]) that: 1) target early exons (first quarter of coding regions); 2) are shared across all relevant transcripts of *rreb1a* and *rreb1b*; 3) have a high “azimuth score”; and 4) have no or very few predicted off-targets with 0-4 mismatches (**ESM Table 2**). As a control gene, we targeted *kita* (ENSDARG00000043317) using a guide RNA designed following the above criteria, which turned out to be identical to CRISPR1-kita (ZDB-CRISPR-180314-3; www.zfin.org) (**ESM Table 2**). *Rreb1a*, *rreb1b* and *kita* (*rreb1a/b* crispants) or *kita* only (controls) were targeted using the Alt-R® CRISPR-Cas9 system (IDT, Belgium) [23] (**ESM Methods**).

#### Imaging of zebrafish larvae

Imaging on zebrafish larvae was performed at 10 days post fertilization (dpf) (**ESM Methods**). Relevant traits for body size (whole body length, dorsal area, lateral area); pancreatic diabetes-related traits (number of insulin-expressing nuclei as a proxy for beta cells; mean and total nuclear volume of insulin-expressing cells; mean and total fluorescence intensity of insulin-expressing nuclei as a proxy of their beta cell insulin expression; islet dimensions); and hepatic diabetes-related traits (liver area; number and size of lipid objects) were quantified in imaging data using custom-written deep-learning algorithms.

#### Glucose and lipid quantification

Imaged larvae or larvae raised to 10 dpf under the same conditions, but collected at 9AM without having been imaged due to time constraints, were stored at −20°C until further processing. Single larvae per well of a 96-well PCR plate were homogenised with a 1.4 mm acid washed zirconium bead (OPS diagnostics, USA) in 88 µL ice cold PBS using a MiniG™ 1600 homogenizer (SPEX® SamplePrep, USA). Samples were centrifuged for 5 minutes at 3500x*g* at 4°C. The supernatant was stored at −80°C until further processing, while the remaining pellet was kept to isolate DNA. Concentrations of glucose, LDL cholesterol, triglycerides and total cholesterol were quantified using enzymatic assays as described previously [24].

#### Identification of control/mutant zebrafish larvae by fragment length analysis and qPCR

Mutant and control larvae were categorised by fragment length analysis (as described by Varshney et al [25]), using DNA extracted from the pellets remaining after homogenization and centrifugation of larvae (**ESM Methods**). Briefly, the remaining pellets were digested using 200 µg/mL proteinase K (ThermoFisher Scientific, Sweden) in 50 μL lysis buffer per larva (10 mM Tris-HCl pH 8, 50 mM KCl, 0.3% Tween 20, 0.3% Igepal, 1 mM EDTA (SigmaAldrich, Sweden)). Samples were incubated at 55°C for 2 hours and at 95°C for 10 minutes, before insoluble particles were removed by centrifugation. *rreb1a* and *rreb1b* amplicons covering the target regions of the guide RNAs were amplified using PCR in separate reactions, using primers (**ESM Table 3**) at a final concentration of 200 nM with either platinum Taq (ThermoFisher Scientific, Sweden) or OneTaq (New England Biolabs, BioNordika Sweden AB, Sweden), following the manufacturer’s protocols. To ascertain how well fragment length analysis quantified CRISPR/Cas9-induced mutagenesis, qPCR was additionally performed in a subset of samples (n=158) using primers described in **ESM Table 4**. For more details, see **ESM Methods**.

### EndoC-βH1 cells

#### Routine cell culture

EndoC-βH1 cells were grown in DMEM, low glucose, pyruvate supplemented with 2% Bovine Serum Albumin Fraction V Fatty acid free (Roche, USA/UK), 50 µM 2-mercaptoethanol, 10 nM nicotinamide (Sigma-Aldrich, USA/UK), 5.5 µg/mL transferrin (Sigma-Aldrich, USA/UK), 6.6 ng/mL sodium selenite (Sigma-Aldrich, USA/UK), 100 U/mL penicillin/streptomycin and 2 mM L-glutamine on cell culture plates coated with DMEM, high glucose supplemented with 1% Extracellular Matrix (Sigma-Aldrich, USA/UK), 2 µg/mL fibronectin (Sigma-Aldrich, USA/UK) and 100 U/mL penicillin/streptomycin at 37°C and 5% CO_2_. Cell culture reagents were manufactured by ThermoFisher Scientific, USA/UK unless otherwise stated. All EndoC-βH1 lines were routinely tested and were negative for mycoplasma.

#### EndoC-βH1 gene silencing

Gene silencing was performed according to the Lipofectamine RNAiMAX^®^ transfection protocol using 25 nM SMART pool ON-TARGETplus siRNAs (PerkinElmer, USA/UK, si*NT*: D-001810-10-05, si*RREB1*: L-019150-00-0005, si*RFX2*: L-011129-00-0005, and si*RFX3*: L-011764-00-0005) diluted in Opti-MEM reduced serum-free medium and 0.4% RNAiMAX^®^ (ThermoFisher Scientific, USA/UK). Silencing efficiency was assessed 96 hours after transfection by qPCR and/or western blot.

#### CRISPR/Cas9-mediated generation of RREB1^KO/KO^ EndoC-βH1 cells

A non-clonal *RREB1*^KO/KO^ EndoC-βH1 cell line was generated using single guide RNAs (sgRNAs) targeting exon 4 (TGACGTCAAGTTCGCCCGC), exon 5 (AGTGCAAATCTTCTCACACA), exon 8 (GTATGGACTGGAGACCCACA), and exon 12 (GACAGACTCCCCCAAAAGCG) of *RREB1* as previously described [26] (see **ESM methods**). EndoC-βH1 cells were transduced at a multiplicity of infection of 10. Selection of transduced cells was performed in 4-6 µg/µL puromycin for seven days. A control EndoC-βH1 line was generated in parallel using a Cas9-only expressing lentivirus.

#### EndoC-βH1 insulin secretion assays

Static insulin secretion assays were performed at 1 mM basal and 20 mM high glucose as previously described [27] (**ESM Methods**). For forskolin-mediated insulin depletion assays, cells were incubated in cell culture media supplemented with 20 mM glucose and 10 µM forskolin for 30 minutes and allowed to recover in 2.8 mM glucose for a further 30 minutes before measuring glucose-stimulated insulin secretion. Insulin secretion was expressed as raw insulin released or as ratio of total secreted insulin to either total insulin content or to cell count on a per-well basis. Values from replicate wells were averaged and normalised to averaged basal insulin secretion of control samples for each experiment to eliminate variability in basal secretion rates across independent experiments.

### Human induced pluripotent stem cells (hiPSC)

#### Routine cell culture

The hiPSC line SB Ad3.1 was obtained from the StemBancc consortium via the Human Biomaterials Resource Centre, University of Birmingham. Human iPSCs were grown in mTeSR1 basal medium supplemented with mTeSR1 5x supplement (StemCell Technologies, UK) and 100 U/mL penicillin/streptomycin on cell culture plates coated with DMEM/F12 (Sigma-Aldrich, UK) supplemented with Matrigel diluted according to the manufacturer’s instructions (hESC qualified, Corning, UK). All lines were maintained at 37°C and 5% CO_2_ and routinely tested negative for mycoplasma.

#### Genome editing of hiPSCs

To generate *RREB1*^WT/WT^ hiPSC clones, two sgRNAs directed to *RREB1* exon 4 (GTCAAGTTCGCCCGCTGGCT) and exon 10 (ACCCCGCGCCAACAGCGGCG) were designed using the MIT CRISPR online design tool (http://crispr.mit.edu) (**ESM Methods**). As the SB hiPSC line is heterozygous for the T2D-protective (p.Asn1171) allele, genome editing was used to generate *RREB1*^WT/WT^ clones with a 141 nucleotide single-stranded oligodeoxynucleotides repair template (Eurogentec, Belgium) containing: 1) the T2D-risk allele (c.3511G, p.Asp1171); 2) a silent mutation to introduce a HincII restriction enzyme site at codon 1170 (c.3510G>C) for genotyping; and 3) a silent mutation in the PAM sequence (c.3507G>C) located in exon 10. The resultant clonal cell lines did not have any of the ten most common coding mutations in the *TP53* gene (**ESM Table 5**) and had normal karyotype, both of which can be affected by the genome editing pipeline.

#### In vitro hiPSC differentiation towards beta-like cells

For differentiation of genome-edited hiPSC lines towards beta-like cells (BLC), hiPSCs were plated in 12-well Corning CellBind plates coated with growth-factor reduced Matrigel (1:30, Corning, UK) at an optimised density of 0.8-1.3×10^6^ cells/well in mTeSR1 supplemented with 10 µM Y-27632 (StemCell Technologies, UK). Once cells had attached (6+ hours), media containing Y-27632 was removed and replaced with mTeSR1. *In vitro* differentiation was started 24 hours after plating following the Rezania et al. protocol [28]. Basal and complete differentiation media can be found in **ESM Table 6**.

#### Flow cytometry

Genome-edited hiPSCs were evaluated for expression of pluripotency markers using the BD Human Pluripotent Stem Cell Transcription Factor Analysis Kit (BD Biosciences, UK). *In vitro* differentiation efficiency was assessed by expression of stage-specific markers of definitive endoderm (CXCR4: 1:40, R and D Systems Cat# FAB173P, RRID: AB_357083) and BLCs (PE Mouse Anti-NKX6.1: 1:40, BD Biosciences Cat# 563023, RRID:AB_2716792; and AF647 Mouse Anti-C-Peptide: 1:200 BD Biosciences Cat# 565831, RRID:AB_2739371). Cells were dissociated into a single-cell suspension, fixed with BD Cytofix^TM^ Fixation Buffer (BD Biosciences, UK) or 4% paraformaldehyde (ThermoFisher Scientific, UK), permeabilised in BD Perm/Wash^TM^ buffer or BD Phosflow Perm Buffer III (BD Biosciences, UK) and stained for cell surface or intracellular markers. Dead cells were excluded using the LIVE/DEAD^TM^ Fixable Violet Dead Cell Stain Kit for 405 nm excitation (ThermoFisher Scientific, UK). Samples were either analysed on the SH800 Cell Sorter (Sony) or the FACSCanto^TM^ II (BD Biosciences, UK). Data analysis was performed using FlowJo^TM^ 10.6.0.

### Gene expression

#### RNA extraction and sequencing

For RNA extraction, cells were lysed in TRIzol® Reagent (ThermoFisher Scientific, UK) and processed following the Direct-zol^TM^ RNA Miniprep kit (Zymo Research, UK) manual. The NEBNext PolyA mRNA Magnetic Isolation Module (New England Biolabs, UK) was used for isolation of polyadenylated transcripts. RNA-Seq libraries were prepared using the NEBNext Ultra Directional RNA Library Kit with 12 cycles of PCR and custom 8 bp indexes (New England Biolabs, UK). Libraries were multiplexed and sequenced on the Illumina HiSeq4000 as 75-nucleotide paired-end reads. Details on RNA-seq analysis can be found in **ESM Methods**.

#### Quantitative RT-PCR

TaqMan® real-time PCR assays were used to measure *RREB1* (Hs00366111_m1), *INS* (H200355773_m1), and *TBP* (H200427620_m1) gene expression. qPCR reactions were performed on a 7900HT Fast Real-Time PCR System (ThermoFisher Scientific, UK) with SDS v2.3 software (Applied Biosystems) and the following conditions: 50°C for 2 minutes, 95°C for 10 minutes, followed by 40 cycles of 95°C for 15 seconds and 60°C for one minute. Cycle thresholds were transformed to gene copy numbers and normalised to the geometric mean of the housekeeping gene *TBP*.

### Protein detection

#### Immunoblotting

Cells were collected using Trypsin-EDTA solution or TrypLE^TM^ Select and lysed in pre-chilled whole cell extraction buffer (20 mM HEPES pH 7.8, 0.42 M NaCl, 0.5% NP-40, 25% glycerol, 0.2 mM EDTA pH 8, 1.5 mM MgCl_2_) supplemented with 1 mM DTT (ThermoFisher Scientific, UK) and 1x protease inhibitor cocktail (Sigma-Aldrich, UK). Protein lysates were quantified using Bradford Assay Reagent, ran on a 4-20% Criterion TGX Stain-Free Precast Gel and transferred to 0.2 µm PVDF membrane (Bio-Rad Laboratories, Inc, UK). Primary antibodies against FLAG (1:10,000, Sigma-Aldrich Cat# F3165, RRID:AB_259529), RREB1 (1:500, Sigma-Aldrich Cat# HPA001756, RRID:AB_1856477 and Atlas Antibodies Cat# HPA034843, RRID:AB_2674357) or β-tubulin (1:2,000, Santa Cruz Biotechnology Cat# sc-365791, RRID:AB_10841919) were used, followed by HRP-conjugated IgG secondary antibodies (1:2,500, ThermoFisher Scientific, UK). Chemiluminescent signals were detected using Clarity Western Enhanced Chemiluminescence Substrate (Bio-Rad Laboratories, Inc, UK) and visualised on a ChemiDoc MP.

#### Immunofluorescence staining

Cells were fixed with 4% paraformaldehyde, permeabilised in 0.001% Triton X-100 (Sigma Aldrich) and blocked in 5% swine-serum. Primary antibodies to RREB1 (1:50, Sigma-Aldrich Cat# HPA001756, RRID:AB_1856477) were incubated at 4°C overnight. The following day, cells were washed before incubation with Alexa Fluor-conjugated secondary antibodies (1:100, ThermoFisher Scientific Cat# A-21206, RRID:AB_2535792 and Cat# A-21435, RRID:AB_2535856) and mounted in Vectashield mounting medium (Vector Laboratories Ltd, UK). Immunostained cells were visualised on a Bio-Rad Radiance 2100 confocal microscope with a 60x 1.0 N.A. water immersion objective. Images were acquired using the LaserSharp 2000 software for three channels (green, red and far-red). For each channel, laser settings were optimised first and the same settings were used for all samples. Image files were exported using the LSM Image Browser 4.2 (Carl Zeiss).

### Human islet studies and genotyping

Donor organs from individuals without type 2 diabetes were obtained with written consent and approval of the Human Research Ethics Board of the University of Alberta (Pro00013094; Pro 00001754). Genotyping was performed on Illumina Omni2.5Exome-8 version 1.3 BeadChip array on DNA extracted from exocrine tissue, spleen, or islets if no other tissue was available. Isolation of human islets and static glucose-stimulated insulin secretion assay were performed as described in the protocols.io repository [29]

### Statistical analysis

Statistical analysis was performed in Prism v8.1.2 (GraphPad Software). Results from multiple experiments are expressed as mean±SEM. A two-tailed unpaired t-test was used to determine p-values for two unmatched groups following a Gaussian distribution. Multiple groups were compared using a two-way ANOVA followed by Sidak’s or Tukey’s multiple comparisons test. Significance was determined using p<0.05. The number of biologically independent experiments (n) are as indicated. Zebrafish data management for fragment length analysis was performed in R. All downstream zebrafish data management and statistical analyses (**ESM Methods**) were performed using Stata MP version 16 (Statacorp, College Station, TX USA).

## Results

### *Rreb1* loss-of-function in zebrafish reduces insulin expression and glucose levels

As the *RREB1* locus is associated with altered diabetes risk and beta cell related traits, we first investigated the impact of loss of *RREB1* at an organismal level. The two orthologues of human RREB1, *rreb1a* and *rreb1b*, were targeted in zebrafish (*Danio rerio*, **ESM Fig. 1a**) at the single cell stage using CRISPR/Cas9 [23]. As the gene structures of *rreb1a* and *rreb1b* are very similar (**ESM Fig. 1a**) and their amino acid sequences are more similar to each other than either is to the human *RREB1* gene (**ESM Fig. 1b**), they are likely remnants of the whole genome duplication in teleost fish [30]. To avoid compensatory effects between the two paralogues, we targeted all relevant transcripts of both genes simultaneously. Survival to 5 dpf was lower in *rreb1a* and *rreb1b* CRISPR-Cas9-targeted crispants than controls, but comparable with crispants for other cardiometabolic candidate genes (**ESM Fig. 1c**).

Using image-based quantification of pancreatic beta cell and hepatic traits, crispants on average had more pancreatic beta cells with a lower average nuclear insulin expression (Fig. 1). Glucose levels were lower in crispants, suggesting a protective effect of mutations in *rreb1a* and *rreb1b*. Crispants were also shorter, had lower LDLc, triglyceride and total cholesterol level, and a smaller liver (Fig. 1). Together, our *in vivo* zebrafish model supports the pancreatic beta cell as a key tissue mediating the association of *RREB1* with disease risk through effects on insulin expression and are consistent with human genetic data regarding effects on size and glycemic traits.

**Figure 1:**
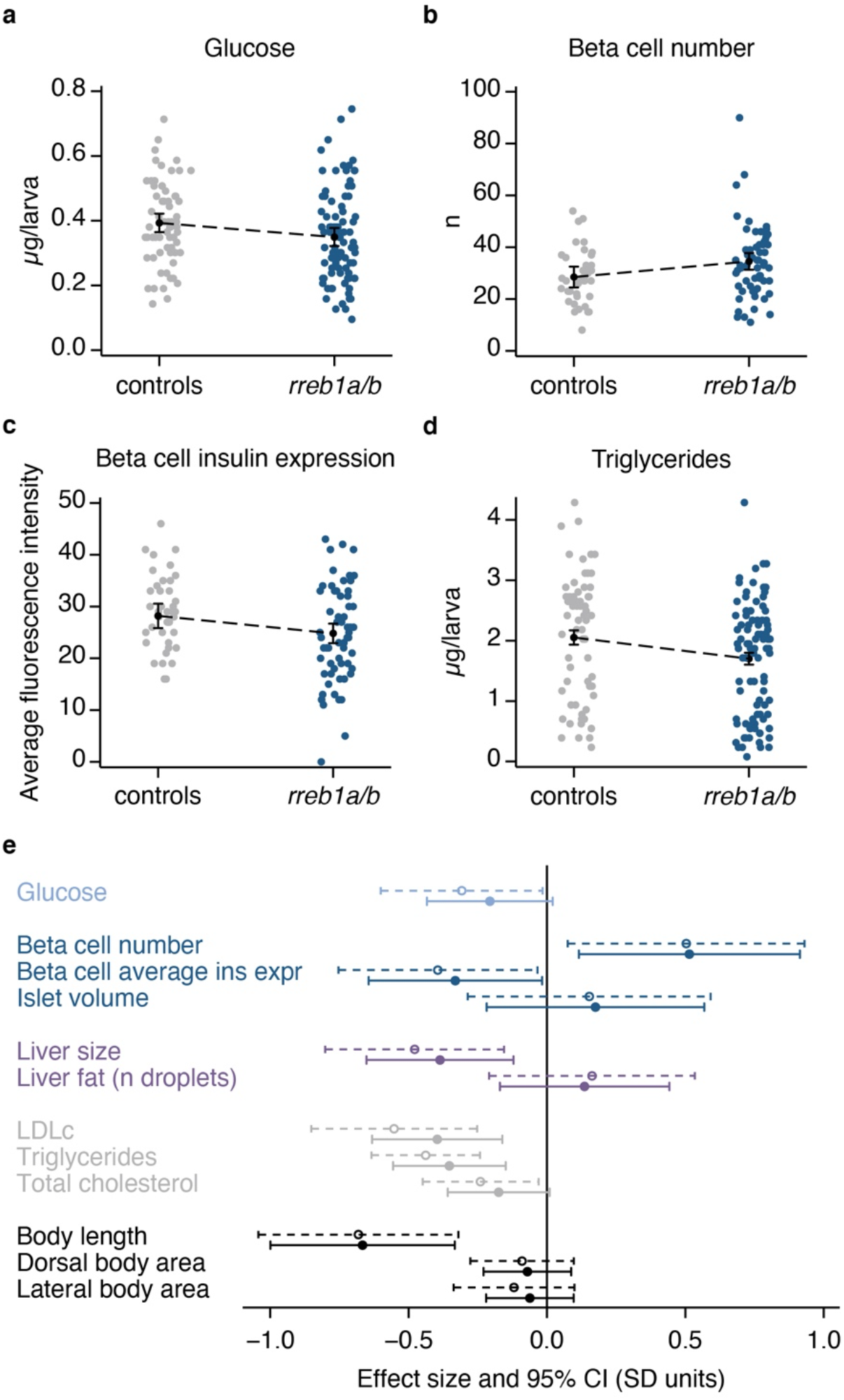
Effect of CRISPR/Cas9-induced mutations in zebrafish *rreb1a* and *rreb1b* on diabetes-related traits. (**a-d**) Individual level data and margins plots for effects of mutations in *rreb1a* and *rreb1b* (vs. sibling controls) on (**a**) glucose levels, (**b**) beta cell number, (**c**) beta cell average insulin expression, and (**d**) triglyceride levels, adjusted for experiment, tank and time of day; and – for glucose and triglyceride levels – imaging (yes/no), and sample position and run. (**e**) Forest plot showing effect sizes and 95% confidence intervals (CI) for 10-day-old CRISPR/Cas9 founders with mutations in *rreb1a/b* and *kita* vs. controls only targeted at *kita*. Dashed 95% CIs (top) reflect results of crispants vs. sibling controls only; full CIs (bottom) show results including 536 additional controls from other experiments performed the same way. All outcomes were inverse normally transformed before linear regression analyses so effect sizes and 95% CIs can be interpreted as z-scores. Adjustments are as described above. Dorsal and lateral body area were additionally adjusted for length.

### RREB1 deficiency reduces *INS* expression and cellular insulin content in human EndoC-βH1

Having established effects on insulin expression following RREB1 loss at an organismal level, we set out to determine whether changes in *RREB1* expression and/or activity altered human beta cell function. To assess the role of RREB1 in a mature beta cell, we performed siRNA knockdown of *RREB1* and assessed glucose-stimulated insulin secretion (GSIS). Transfection of siRNAs in EndoC-βH1 cells reduced expression of *RREB1* by 34±9% (Fig. 2a), *INS* transcript levels by 16±4% (Fig. 2b), and cellular insulin content by 32±11% (Fig. 2c) compared with siNT controls. si*RREB1* knockdown cells had similar insulin secretion, either raw (**ESM Fig. 2a**) or normalised for insulin content (**ESM Fig. 2b**). As loss of *RREB1* decreased insulin content, we normalised insulin secretion for cell count (**ESM Fig. 2c**). There was no effect on basal (1 mM) or glucose-stimulated (20 mM) insulin secretion following *RREB1* knockdown (Fig. 2d).

**Figure 2:**
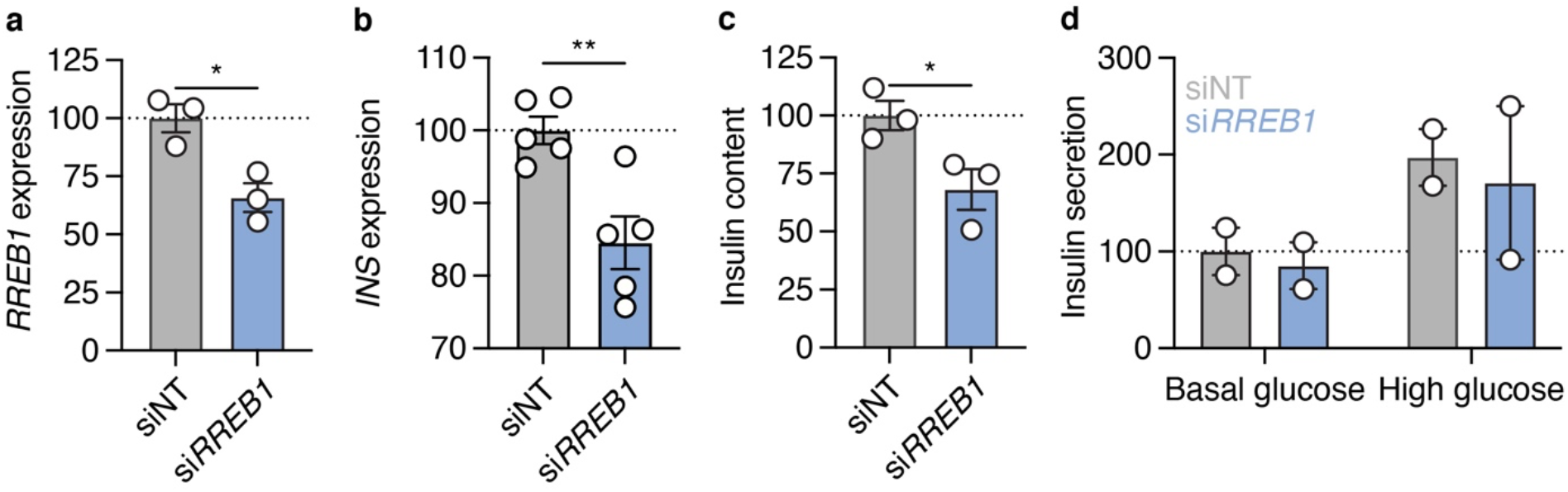
Partial loss of RREB1 reduces cellular insulin content in mature beta cells. (**a-b**) Gene expression (% of siNT, normalised to housekeeping gene) of (**a**) *RREB1* and (**b**) *INS* in si*RREB1* knockdown and siNT control EndoC-βH1 cells. (**c**) Cellular insulin content (% of siNT) and (**d**) glucose stimulated insulin secretion (% of siNT at basal glucose, normalised to cell count) measured in siNT and si*RREB1* EndoC-βH1 cells. Data are presented as mean±SEM. \**p* <0.05, \*\**p* <0.01. (Unpaired t-test)

As we previously identified phenotypic differences between transient and long-term loss-of-function in the EndoC-βH1 model [26], we next used CRISPR-Cas9 to generate pooled knockout *RREB1* EndoC-βH1 cells. To control for the genome editing pipeline, *RREB1*^WT/WT^ cells were generated to express Cas9 protein without sgRNAs targeting the genome. Four sgRNAs targeting the protein coding sequence of *RREB1* were used to generate *RREB1*^KO/KO^ EndoC-βH1 cells (**ESM Fig. 3a**). *RREB1*^KO/KO^ cells had a near complete loss of RREB1 protein compared to the parental and *RREB1*^WT/WT^ cells (**ESM Fig. 3b**). CRISPR-Cas9 mediated loss of RREB1 in EndoC-βH1 cells resulted in a 35±14% reduction in *INS* expression (Fig. 3a) and a 44±7% reduction in cellular insulin content (Fig. 3b) compared to *RREB1*^WT/WT^ cells. Similar to transient knockdown, insulin secretion from *RREB1*^KO/KO^ EndoC-βH1 cells was not significantly different from control *RREB1*^WT/WT^ cells at basal or high glucose when normalised for cell count (Fig. 3c). Taken together, loss of *RREB1* in human beta cells reduces cellular insulin content but does not affect glucose-stimulated insulin secretion.

**Figure 3:**
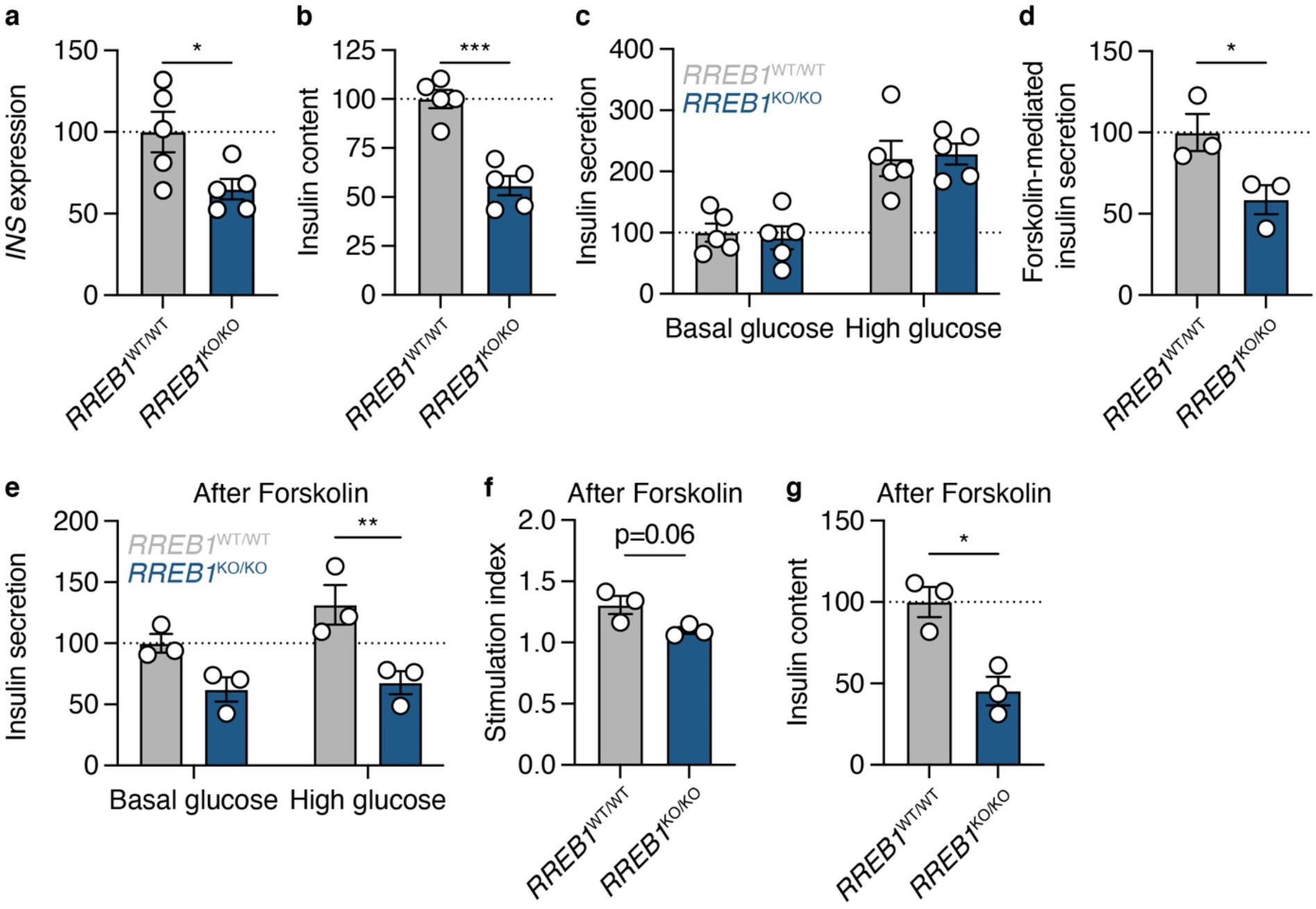
RREB1 knockout reduces cellular insulin content in mature beta cells. (**a-c**) *RREB1*^KO/KO^ EndoC-βH1 cells were assessed for (**a**) *INS* gene expression (% of *RREB1*^WT/WT^, normalised to housekeeping gene), (**b**) cellular insulin content (% of *RREB1*^WT/WT^), and (**c**) glucose stimulated insulin secretion (% of *RREB1*^WT/WT^ at basal glucose, normalised to cell count). (**d**) Forskolin-mediated insulin secretion at high glucose (% of *RREB1*^WT/WT^, normalised to cell count). (**e-g**) After stimulation with forskolin, (**e**) glucose stimulated insulin secretion (% of *RREB1*^WT/WT^ at basal glucose, normalised to cell count), (**f**) stimulation index, and (**g**) insulin content (% of *RREB1*^WT/WT^, normalised to cell count) were measured. Data are presented as mean±SEM. \**p* <0.05, \*\**p* <0.01, *p* ***<0.001. (Unpaired t-test or two-way ANOVA followed by Sidak’s multiple comparisons test)

To assess the effect of lower insulin availability in *RREB1*^KO/KO^ beta cells under conditions of prolonged insulin demand, cells were stimulated with 20 mM glucose in combination with the cAMP-elevating agent forskolin (10 μM) for 30 minutes prior to evaluation of GSIS. *RREB1*^KO/KO^ EndoC-βH1 cells secreted 41±15% less insulin in response to forskolin stimulation compared to control cells (Fig. 3d). Assessment of insulin release after forskolin-mediated docked granule depletion showed significantly reduced insulin secretion in response to a glucose stimulus in RREB1-deficient EndoC-βH1 cells (Fig. 3e). While the response to glucose after the forskolin challenge was blunted in both *RREB1*^KO/KO^ and control EndoC-βH1 cells (stimulation index: *RREB1*^KO/KO^ 1.1±0.1; *RREB1*^WT/WT^ 1.3±0.1), RREB1-depleted cells tended to recover more poorly than control EndoC-βH1 cells (p=0.06) (Fig. 3f). After forskolin treatment, the insulin content was reduced 55±13% in *RREB1*^KO/KO^ EndoC-βH1 cells (Fig. 3g), suggesting that loss of RREB1 negatively impacts insulin secretion during periods of prolonged demand.

### RREB1 is a novel transcriptional activator and repressor in mature beta cells

As RREB1 is a transcription factor, we next performed transcriptomic analysis in EndoC-βH1 cells following siRNA-mediated knockdown and CRISPR-Cas9 knockout. In total, 2,144 differentially expressed genes (DEGs) were detected between siNT and si*RREB1* treated samples, with slightly more upregulated genes (56%) in the RREB1-depleted cells (**ESM Table 7**). Roughly half (55% and 56% of upregulated and downregulated, respectively) of the DEGs corresponded to predicted RREB1 target genes identified in JASPAR and TRANSFAC databases [31, 32]. Enriched biological terms and pathways among all upregulated DEGs included processes associated with neurons, such as ‘nervous system development’, ‘neuronal system’, ‘synaptic signalling’, and ‘axon guidance’, likely reflecting the phenotypic and transcriptomic similarities between neurons and beta cells [33]. In addition, terms relating to exocytotic processes, like ‘regulation of exocytosis’, ‘synaptic vesicle exocytosis’, and ‘transmission across chemical synapses’ were also enriched in upregulated DEGs, consistent with the role of RREB1-regulated genes in insulin secretion.

Differential gene expression analysis identified 2,604 DEGs between *RREB1*^KO/KO^ and *RREB1*^WT/WT^ EndoC-βH1 cells with more than half (66%) being up-regulated as a consequence of RREB1 loss (**ESM Table 8**). *RREB1* gene expression was elevated in *RREB1*^KO/KO^ cells compared to wildtype cells (q=2.84×10^−21^, log_2_FC=0.7444). However, increased expression was due to exons in the 5’-UTR (q<0.001) that were not targeted by the four sgRNAs, consistent with a genetic compensation for loss of RREB1. Other upregulated genes included transcripts involved in insulin secretion and processing (*GCK*, *CHGB*, *PCSK1*, *SNAP25*, *SCG2*), voltage-sensitive Ca^2+^ channel subunits (*CACAN1A*, *CACNA1B*, *CACAN1C*, *CACAN1D*, *CACAN1E*) and cell-to-cell communication (*GJD2*, *NCAM1*, *PTPRN*), suggesting a potential compensatory effect for the reduced insulin content (**ESM Table 8**). Accordingly, upregulated DEGs were enriched for biological terms related to exocytosis and insulin secretion (**ESM Table 9**). Expression of *NEUROD1*, which encodes a well-established regulator of the *INS* gene [34], was significantly downregulated in *RREB1*^KO/KO^ compared to *RREB1*^WT/WT^ EndoC-βH1 cells. Finally, a subset of DEGs encode transcription factors that are important for stem cell fate (*NANOG*, *KLF4*) or early endoderm formation (*HHEX*, *GATA4* and *GATA6*), suggesting that RREB1 may regulate genes involved in endocrine cell differentiation.

### RREB1 loss-of-function during *in vitro* differentiation affects endocrine progenitor development

To address the role of RREB1 during endocrine cell differentiation, we generated multiple isogenic *RREB1*^WT/WT^ and *RREB1*^KO/KO^ hiPSC lines. Four independent *RREB1*^KO/KO^ lines were generated using two sgRNAs that target sequences either close to the start codon (exon 4) or in a distal exon (exon 10). Both sgRNAs are located in genomic regions that are common to all protein-coding *RREB1* transcripts and generated a ∼50 kb deletion (**ESM Fig. 4a**). As sequencing of the SB Ad3.1 hiPSC line revealed heterozygosity for the common T2D-associated variant rs9379084 (c.3511G>A, p.Asp1171Asn) in *RREB1*, *RREB1*^WT/WT^ lines were genetically edited to be homozygous for the major allele at rs9379084, associated with higher risk of T2D (c.3511G, p.Asp1171Asp) (**ESM Fig. 4a**). Quantification of RREB1 protein showed no difference in expression levels between the three edited *RREB1*^WT/WT^ clones and an unedited, parental SB Ad3.1 (p.Asp1171Asn) hiPSC line (**ESM Fig. 4b-c**). RREB1 protein was not detectable in any of the four *RREB1*^KO/KO^ hiPSC lines by western blot and immunofluorescent staining (**ESM Fig. 4b-d**). All gene-edited *RREB1* hiPSC lines expressed pluripotency markers (OCT4, SOX2, NANOG and SSEA-4) (**ESM Fig. 4e**), had typical hiPSC morphology, a diploid karyotype, and had not acquired any of the 10 most frequently detected coding mutations in *TP53* from the genome editing process (**ESM Table 5**).

To model endocrine pancreas development, we differentiated the *RREB1*^WT/WT^ and *RREB1*^KO/KO^ hiPSC lines along the endocrine lineage into beta-like cells [28] and performed transcriptomic analysis at all seven stages of *in vitro* differentiation (Fig. 4a). *RREB1* was expressed at all stages of beta cell differentiation in *RREB1*^WT/WT^ cells and its expression was significantly reduced in *RREB1*^KO/KO^ lines (Fig. 4b). Stage-specific marker expression revealed that *RREB1*^KO/KO^ and *RREB1*^WT/WT^ lines followed established endocrine development and generated beta-like cells characterised by co-expression of NKX6.1 and C-peptide (**ESM Fig. 5a-b**). Principal component analysis revealed that both *RREB1*^WT/WT^ and *RREB1*^KO/KO^ samples clustered by developmental stage in the expected pattern, with more variability observed in the later stages (Fig. 4c). Differential expression analysis at each differentiation stage revealed that loss of RREB1 resulted in a total of 5,476 DEGs between *RREB1*^WT/WT^ and *RREB1*^KO/KO^ lines, of which 159 were common to all developmental stages (**ESM Table 10**). The majority of DEGs were upregulated in the *RREB1*^KO/KO^ lines (63±5%) and found at the endocrine progenitor stage. Upregulated DEGs in *RREB1*^KO/KO^ hiPSC-derived endocrine progenitor and endocrine cells were enriched for genes involved in the ‘regulation of gene expression in endocrine committed (NEUROG3+) progenitor cells’ and ‘insulin secretion’ or ‘regulation of insulin secretion’, respectively (**ESM Table 11**). Interestingly, transcript expression of the endocrine progenitor marker *NEUROG3* was significantly higher in *RREB1*^KO/KO^ hiPSC-derived endocrine progenitor cells (**ESM Fig. 5c**), suggesting accelerated differentiation towards the endocrine lineage.

**Figure 4:**
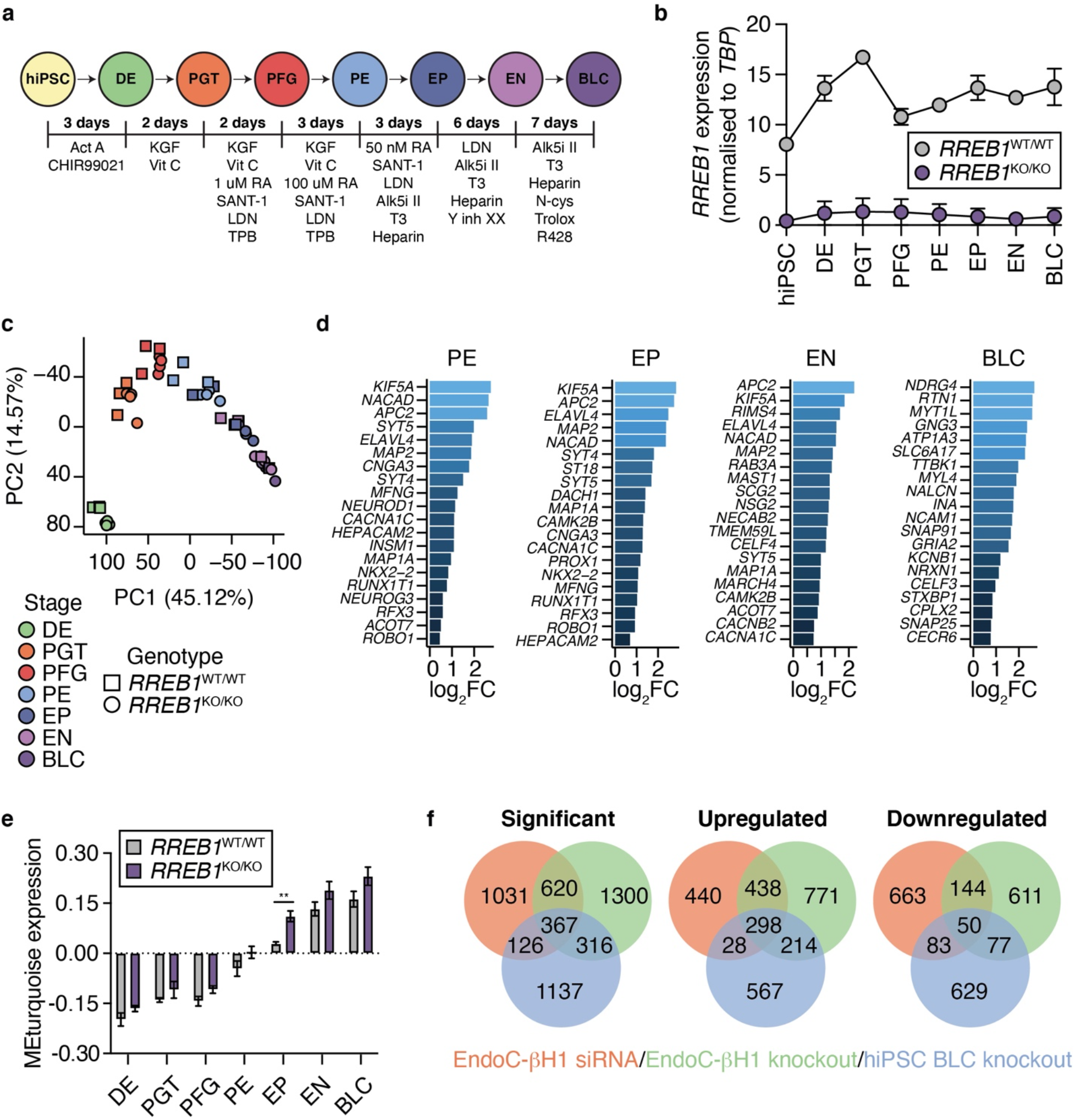
Transcriptomic analysis reveals altered expression of pro-endocrine genes following loss of *RREB1* in human beta cells. (**a**) Schematic of *in vitro* differentiation protocol stages: human induced pluripotent stem cell (hiPSC), definitive endoderm (DE), primitive gut tube (PGT), posterior foregut (PFG), pancreatic endoderm (PE), endocrine progenitor (EP), endocrine (EN) and beta-like cells (BLC). Growth factors and small molecules (listed underneath) were added for the indicated amount of time. (**b**) *RREB1* expression in *RREB1*^WT/WT^ and *RREB1*^KO/KO^ hiPSC lines during *in vitro* differentiation toward BLC. (**c**) The first two principal components (PC1, PC2) were calculated using normalised gene counts of *RREB1*^KO/KO^ (circles) and *RREB1^WT^*^/WT^ (squares) cell lines for all seven stages of *in vitro* beta cell differentiation. (**d**) Differential expression of endocrine cell genes in PE, EP, EN, and BLCs. (**e**) Analysis of modules of co-expressed genes using WGCNA bar plot showing module epigengene (ME) expression of the module enriched for endocrine progenitor and endocrine genes. (**f**) Venn diagrams of the overlap of DEGs between si*RREB1* knockdown EndoC-βH1, *RREB1*^KO/KO^ EndoC-βH1, and *RREB1*^KO/KO^ hiPSC-derived BLCs. n=3-4. Data are presented as mean±SEM. \*\**p* <0.01.

Using stage-specific markers identified in human fetal pancreata [35], hypergeometric enrichment analyses revealed an enrichment of endocrine progenitor markers (*NEUROG3*, *NEUROD1*, *NKX2.2*, *RFX3*, *CACNA1C*) among genes upregulated in *RREB1*^KO/KO^ lines in pancreatic endoderm (q=4.0×10^−83^), endocrine precursor (q=6.3×10^−104^) and endocrine-like (q=5.5×10^−43^) cells (Fig. 4d). Among genes upregulated in *RREB1*^KO/KO^ beta-like cells, there was an enrichment of genes implicated in insulin exocytosis (*SNAP25*, *STXBP1*, *NRXN1*) (q=1.6×10^−35^) (Fig. 4d). Downregulated DEGs were enriched in early and late pancreatic progenitors (q=3.2×10^−28^ and q=7.2×10^−24^, respectively), including two acinar cell markers (*CPA2* and *NR5A2*), the multipotent pancreatic progenitor transcription factor *HNF1B* [36, 37], and members of the Notch signalling (*NOTCH1*, *NOTCH2*, *JAG1*) and EGF and FGF pathways (*ERBB3*, *FGFR2*). To identify co-expressed genes that may be regulated by RREB1, weighted gene co-expression network analysis (WGCNA) [38, 39] was performed. The module eigengene turquoise (MEturquoise) enriched for endocrine progenitor and endocrine genes (q=4.56×10^−18^ and q=3.98×10^−19^, respectively), showed significant expression differences between *RREB1*^WT/WT^ and *RREB1*^KO/KO^ clones (Fig. 4e; **ESM Table 12**). Interestingly, a subset of significantly upregulated and downregulated genes were shared amongst the EndoC-βH1 siRNA, EndoC-βH1 *RREB1*^KO/KO^ and hiPSC-derived *RREB1*^KO/KO^ BLCs (Fig. 4f), suggesting a common RREB1 regulatory network between developing and mature beta cells.

### Loss of RREB1 increases RFX motif activity during endocrine cell differentiation and in mature beta cells

Computational prediction of upstream regulators of the DEGs in hiPSC-derived BLCs and EndoC-βH1 cells using iRegulon [40] highlighted RREB1, as well as the RFX transcription factor family (Fig. 5a). The RFX family is comprised of eight members and is characterised by a highly conserved DNA binding domain [41, 42]. Loss of *RREB1* significantly increased *RFX2* expression (q=6.47×10^−5^, log_2_FC=0.3629) and decreased *RFX6* (p=3.06×10^−8^, log_2_FC=-0.4188) in EndoC-βH1 cells (Fig. 5b-c), while expression of *RFX3* was unchanged (**ESM Fig. 6**). Interestingly, while RFX2 protein expression was markedly increased in *RREB1*^KO/KO^ EndoC-βH1 cells (22.26±0.10-fold, p=0.0123), loss of RREB1 did not affect RFX6 protein expression in mature beta cells (Fig. 5d-e).

**Figure 5:**
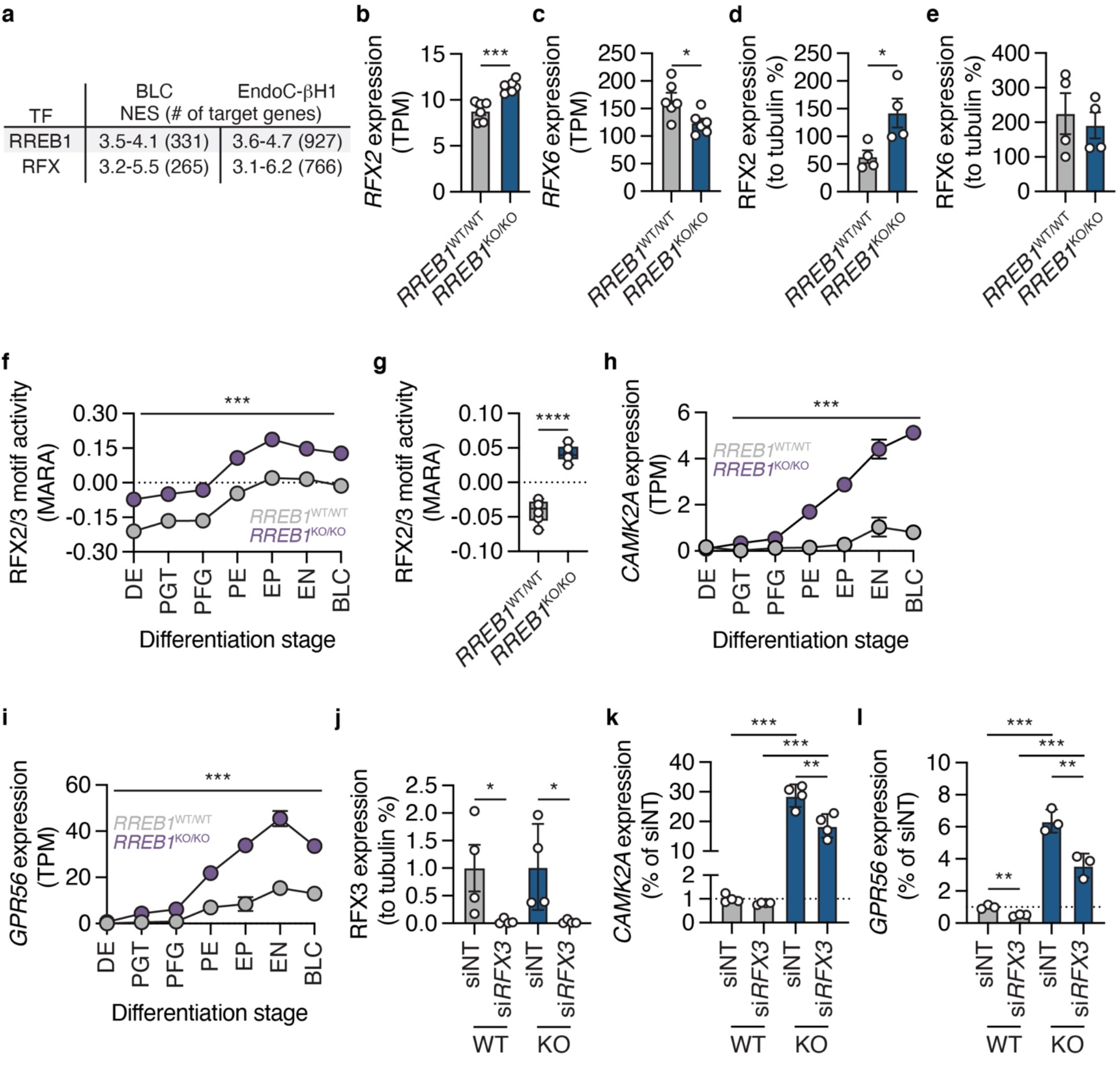
RREB1 deficiency affects RFX motif activity. (**a**) Most common TF motifs in hiPSC-derived BLCs and EndoC-βH1 cells of 9,713 position weight matrices (PWMs) and 1,120 ENCODE ChIP-Seq tracks (centred 10kb around TSS) tested. NES, normalised enrichment score with cutoff set to >3 (corresponding to false discovery rate (FDR) of 3-9%); # targets, number of targets for TF motif with highest NES; RFX comprises RFX1-6, 8, RFXANK and RFXAP; TF, transcription factor. (**b-c**) Expression of (**b**) *RFX2* and (**c**) *RFX6* mRNA levels in *RREB1*^WT/WT^ and *RREB1*^KO/KO^ EndoC-βH1 cells. (**d-e**) Protein quantification of (**d**) RFX2 and (**e**) RFX6 in RREB1-deficient cells. (**f**) RFX2/3 motif activity profiles for *RREB1*^KO/KO^ and *RREB1^WT^*^/WT^ lines during *in vitro* differentiation towards BLCs calculated using MARA. (**g**) RFX2/3 motif activities in *RREB1*^WT/WT^ and *RREB1*^KO/KO^ EndoC-βH1 cells. (**h-i**) Expression of RFX2/3 target genes (**h**) *CAMK2A* and (**i**) *GPR56* during *in vitro* differentiation of *RREB1*^WT/WT^ *RREB1*^KO/KO^ lines. (**j**) RFX3 protein quantification (% of siNT, normalised to tubulin) in *RREB1*^WT/WT^ and *RREB1*^KO/KO^ EndoC-βH1 cells following siNT and si*RFX3* transfection. (**k-l**) Gene expression of (**k**) *CAMK2A* and (**l**) *GPR56* (% of siNT, normalised to housekeeping gene) in *RREB1*^WT/WT^ and *RREB1*^KO/KO^ EndoC-βH1 cells following siRNA-mediated deletion of *RFX3*. n=3-6. Data are presented as mean±SEM. *p <0.05, **p <0.01, ***p <0.001, ****p <0.0001. (Unpaired t-test)

Motif activity response analysis (MARA), a further approach to predict genome-wide regulatory interactions that underlay gene expression variation across *RREB1*^KO/KO^ and *RREB1*^WT/WT^ clones, predicted RFX2 and RFX3 as key transcription factors driving differential gene expression across *RREB1*^KO/KO^ and *RREB1*^WT/WT^ cells during beta cell differentiation (RFX2/3 Z=12.43) (Fig. 5f) and in EndoC-βH1 cells (RFX2/3 Z=8.45) (Fig. 5g). RFX2/3 target genes *CAMK2A* [43] and *GPR56* [44] were among the differentially expressed genes showing the strongest upregulation in *RREB1*^KO/KO^ clones across all seven differentiation stages (**Fig. h-i**). Taken together, the transcriptomic analysis revealed *RFX* family members as potential targets of RREB1 in beta cells.

Our *in silico* approaches were unable to distinguish between RFX2 and RFX3 owing to their similar binding motif. Thus, we next used RNA interference mediated inhibition of both *RFX2* and *RFX3* to determine if the changes in gene expression following RREB1 loss could be mirrored by modulating RFX proteins in beta cells. Loss of RFX2 protein following RNA interference (**ESM Fig. 6b**) did not impact expression of target genes *CAMK2A* and *GPR56* (**ESM Fig. 6c-d**). RNA interference driven reductions in RFX3 protein levels (Fig, 5j) partially rescued increased *CAMK2A* expression in *RREB1*^KO/KO^ beta cells (Fig. 5k). *GPR56*, an established RFX3 target gene, was downregulated in both *RREB1^KO/KO^*(44±10%) and *RREB1^WT/WT^* (51±5%) beta cells as a consequence of RFX3 depletion (Fig. 5l), strengthening RFX3 as transcriptional regulator affected by loss of RREB1 in mature beta cells.

### Carriers of T2D-risk alleles in RREB1 have altered beta cell function

RREB1 loss-of-function in a human beta cell model negatively impacted insulin content and secretion. To determine whether all three independent signals at the *RREB1* locus influence pancreatic islet function, we quantified glucose-stimulated insulin secretion of *ex vivo* human islets stratified by genotype. For the causal coding variant (rs9379084), glucose-stimulated insulin secretion was paradoxically higher in carriers of the T2D-risk allele (G; p.D1171); although insulin content was reduced, it did not reach statistical significance (Fig. 6a-b). Neither of the index variants at the two regulatory signals (rs9505097 and rs112498319) influenced insulin content (Fig. 6c and e). However, carriers of the rs112498319 T2D-risk allele (C) on average had lower glucose-stimulated insulin secretion (Fig. 6f). Together, these results support a role for RREB1 in human pancreatic islet function and suggest that at least 2 of the 3 signals at the locus alter islet-cell function.

**Figure 6:**
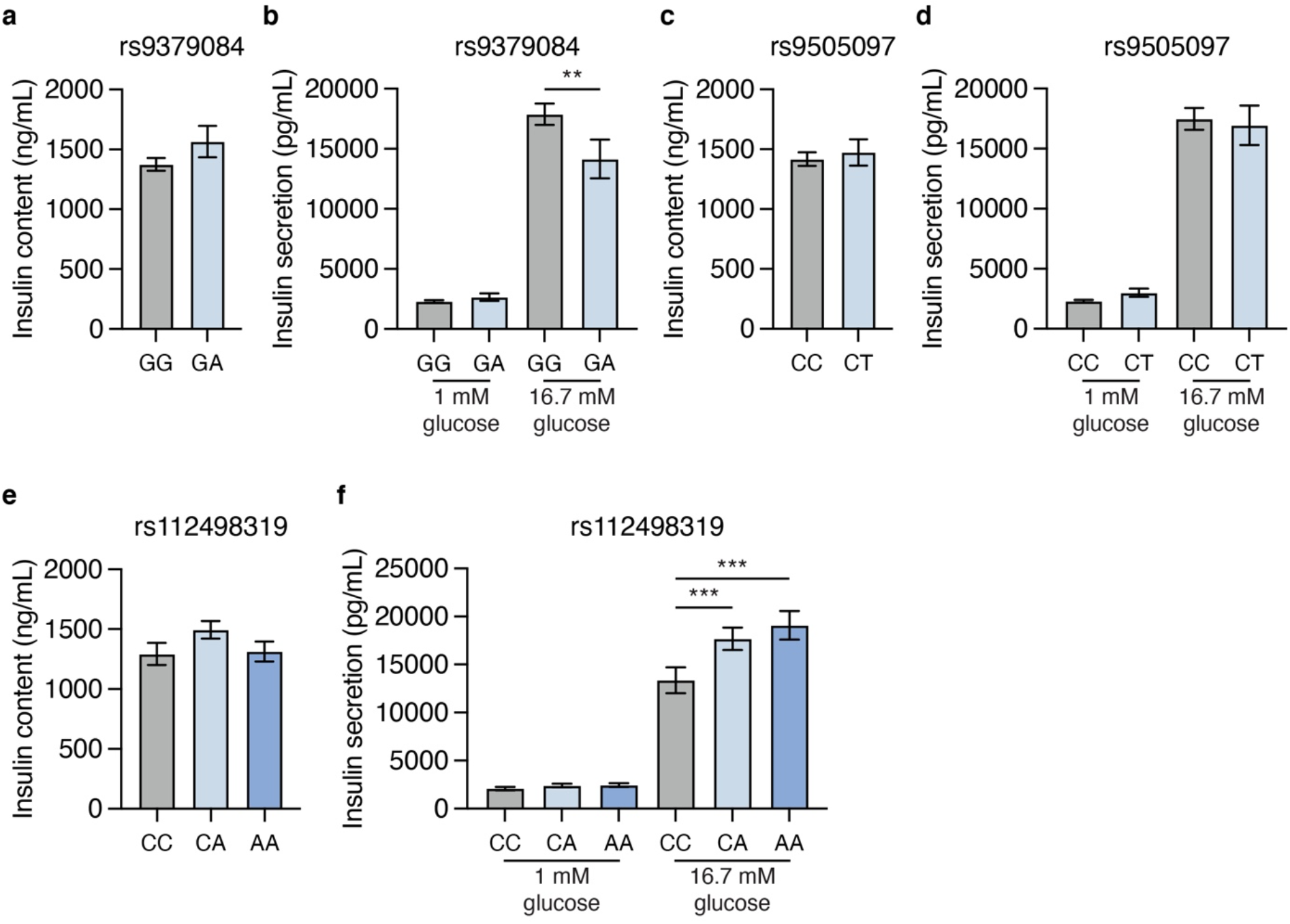
Genetic variation at the *RREB1* locus influences human beta cell function. Insulin content and glucose-stimulated insulin secretion of human donor islets from carriers of *RREB1* (**a-b**) rs9379084, (**c-d**) rs9505097, and (**e-f**) rs112498319 variants. (Mann-Whitney test or one-way ANOVA)

## Discussion

Our understanding of the genetic landscape of T2D has increased substantially [2, 5, 45] and current efforts are focused on translating these genetic discoveries into disease mechanisms. Here, we characterised the role of the T2D-associated gene *RREB1* in beta cell development and function. Our *in vivo* zebrafish model lacking *rreb1a* and *rreb1b* had reductions in beta cell insulin expression. Loss of RREB1 reduced insulin gene expression and cellular insulin content in EndoC-βH1 cells, resulting in impaired glucose stimulated insulin secretion under prolonged stimulation. Transcriptomic analysis identified RREB1 as a novel transcriptional activator and repressor in developing and mature human beta cells. Isolated human islets from carriers of the *RREB1* coding allele that reduces diabetes risk (p.Asn1171) had lower glucose-stimulated insulin secretion. Taken together, our data are consistent with T2D-protective alleles in RREB1 resulting in a loss-of-function. The contradictory finding that carriers of the *RREB1* protective allele have lower insulin secretion but are protected from T2D, hints at potential additional functions of RREB1 in other diabetes relevant tissues (e.g. insulin responsive tissues).

Loss of RREB1 led to a significant increase in transcriptional activity of *RFX2* and *RFX3* in both the hiPSC-based developmental and the mature EndoC-βH1 model. In line with this, RFX target genes *GPR56* and *CAMK2A* were significantly upregulated in *RREB1*^KO/KO^ cell models. While RFX3 and RFX6 have been implicated in beta cell development, formation and function [46–51], a role for RFX2 has not yet been described. Human mapping of protein-protein interactions revealed that RFX6 physically interacts with RFX2 and RFX3 [52]; however, whether RFX2 and RFX3 form heterodimers in beta cells to cooperatively regulate gene expression is currently unknown. Loss of RREB1 in mature beta cells increased the expression of RFX2 transcript and protein, highlighting RREB1 as a transcriptional repressor of *RFX2* in mature beta cells and likely during endocrine cell differentiation. A study aimed at the prediction of upstream transcriptional regulators of RFX genes using transcription factor binding profile analysis did not identify the RREB1 transcription factor binding site as being statistically over-represented in *RFX* promoters [42], suggesting that RREB1 regulation of RFX expression is likely to be indirect.

One of the consistent phenotypes across the human and zebrafish models is the reduction in insulin expression. An important transcriptional activator of the *INS* gene is the transcription factor NEUROD1 [34], which has been shown to co-occupy the *Ins1* and *Ins2* promoters with the CtBP/RREB1 co-repressor complex in murine beta cells [53]. As such, it is tempting to hypothesize that RREB1 and NEUROD1 may also interact to transcriptionally regulate expression of the human *INS* gene.

In the absence of a validated assay to quantify zebrafish insulin protein levels, we used a transgenically expressed beta cell reporter with H2B-mCherry expression under the control of the insulin promoter. Downsides of this approach are that insulin promoter activity may not reflect the more physiologically relevant plasma insulin concentration, and that transcriptional regulation and H2B-mCherry turnover may not reflect endogenous insulin expression. However, the integration over a longer time frame, as happens with a reporter like H2B-mCherry, could be considered advantageous, as it is less prone to short term effects introduced by interindividual differences in nutritional status. Moreover, we previously observed lower H2B-mCherry expression upon exposure to 3% glucose from day 5 to 10 post-fertilisation, or to mutations in e.g. *pdx1*, suggesting that this readout can provide valid insights.

Our study - in which we characterised an *in vitro* zebrafish model, two complementary *RREB1*^KO/KO^ human cellular beta cell models, and *ex vivo* islet cell function from human carriers of *RREB1* alleles - strongly suggests a novel role for RREB1 in beta cell development and function through a transcriptional effect of RREB1 on endocrine cell-specific gene expression. Identification of RREB1 as regulator of multiple genes of known importance in endocrine cell development and insulin secretion has important implications for future evaluation of type 2 diabetes risk-associated variants, as they might exert their effects through modification of RREB1 binding sites in islet cells.

## Supporting information

ESM Methods and Figures

ESM Table 6

ESM Table 7

ESM Table 8

ESM Table 9

ESM Table 10

ESM Table 11

ESM Table 12

## Abbreviations

BLC: beta-like cells
CI: confidence interval
DE: definitive endoderm
DEG: differentially expressed genes
dpf: day(s) post fertilization
EP: endocrine progenitor
EN: endocrine cells
FDR: false discovery rate
GSIS: glucose-stimulated insulin secretion
hiPSC: human induced pluripotent stem cell
KO: knockout
NES: normalised enrichment score
PE: pancreatic endoderm
PFG: posterior foregut
PGT: primitive gut tube
PWM: position weight matrices
RREB1: Ras-responsive element binding protein 1
sgRNA: single guide RNA
T2D: type 2 diabetes
TPM: transcript per million
WGCNA: weighted gene co-expression network analysis
WT: wildtype

## Acknowledgements

The authors thank Dr Toryn Poolman and Daniela Moralli (University of Oxford) for technical support. For the zebrafish studies, we would like to acknowledge Anastasia Emmanouilidou for methods development and project management; Hanqing Zhang and Amin Allalou for image analysis; João Campos-Costa for line maintenance; Ghazal Alavioon for help with micro-injections; and Kanaka Lakshmi Gurrala for zebrafish care taking. pLentiCRISPRv2 and pX330-U6-Chimeric_BB-CBh-hSpCas9 were provided by Dr Feng Zhang. We thank the Human Organ Procurement and Exchange (HOPE) program and Trillium Gift of Life Network (TGLN) for their work in procuring human donor pancreas for research, and James Lyon (Edmonton) and Nancy Smith (Edmonton) for their work in human islet isolation at the Alberta Diabetes Institute IsletCore (www.isletcore.ca). We especially thank the organ donors and their families for their kind gift in support of diabetes research.

## Data Availability

The datasets generated during and/or analysed during the current study are available in the European Genome Archive repository https://ega-archive.org under accession number EGAS00001006314.

## Authors’ relationships and activities

AKR hold stock options in AstraZeneca. As of January 2020, AM is an employee of Genentech, and a holder of Roche stock. ALG discloses that her spouse is an employee of Genentech and hold stock options in Roche. All other authors declare no interests that could be considered conflicting.

## Contribution statement

KKM, NAJK, and FA designed experiments, and interpreted the data (human studies); CM and MdH designed experiments and interpreted the data (zebrafish studies); KKM, NAJK, CM, FA, AFS, AKR, AB, EM, JEMF, JMT, AM, and AL conducted experiments and collected data; KKM, NAJK, CM, JMT, MdH, and PEM analysed data; HS, MPA, and AWA provided analytical support; GY, AM, and PEM contributed to discussion; ALG developed the concept; ALG and BD supervised the project; KKM, NAJK, CM, MdH, and ALG wrote the manuscript with all co-authors approving the final manuscript. ALG is the guarantor of this work and, as such, has full access to all the data in the study and takes responsibility for the integrity of the data and the accuracy of the data analysis.

## ESM Tables

ESM Table 1: RREB1 orthologues in Danio rerio (Ensembl release 105 - Dec 2021).

ESM Table 2: Gene IDs, CRISPOR scores and gRNA target sequences for zebrafish genes *rreb1a*, *rreb1b* and *kita*.

ESM Table 3: Sequences of primers used for fragment length analysis.

ESM Table 4: Sequences of primers used for qPCR based analysis of mutagenesis efficiency.

ESM Table 5: Ten most common coding mutations in *TP53* that were assessed in genome-edited hiPSC lines.

ESM Table 6: Composition of basal and complete differentiation media.

ESM Table 7: List of DEGs between si*RREB1* and siNT EndoC-βH1 cells.

ESM Table 8: List of DEGs between *RREB1*^KO/KO^ and *RREB1*^WT/WT^ EndoC-βH1 cells.

ESM Table 9: Enriched biological terms and pathways among DEGs between *RREB1*^KO/KO^ and *RREB1*^WT/WT^ EndoC-βH1 cells.

ESM Table 10: List of DEGs between *RREB1*^KO/KO^ and *RREB1*^WT/WT^ hiPSC during seven stages of *in vitro* beta cell differentiation.

ESM Table 11: Enriched biological terms and pathways among DEGs up-regulated in *RREB1*^KO/KO^ lines at seven distinct stages of *in vitro* beta cell differentiation.

ESM Table 12: Module eigengenes identified in WGCNA for *RREB1*^KO/KO^ and *RREB1*^WT/WT^ lines during seven stages of *in vitro* beta cell differentiation.

## ESM Figure Legends

ESM Fig. 1: Characterisation of zebrafish model.

ESM Fig. 2: Characterisation of glucose-stimulated insulin secretion from si*RREB1* EndoC-βH1 cells.

ESM Fig. 3: Generation and characterisation of *RREB1*^KO/KO^ EndoC-βH1 cells.

ESM Fig. 4: Generation and characterisation of *RREB1*^KO/KO^ hiPSC lines.

ESM Fig. 5: Characterisation of *in vitro* differentiation of *RREB1*^KO/KO^ and *RREB1*^WT/WT^ clones towards beta-like cells

ESM Fig. 6: RREB1 deficiency in EndoC-βH1 cells alters gene expression of *RFX* family members.

