## Supplementary material for "Loss of RREB1 in pancreatic beta cells reduces cellular insulin content and affects endocrine cell gene expression": ESM Methods and Figures

**Electronic Supplementary Material**

**ESM Methods**

*CRISPR/Cas9 in zebrafish*

crRNA with the selected target sequences and Alt-R® CRISPR-Cas9 tracrRNA (IDT, 072533) were resuspended at 100 µM in Duplex Buffer and stored at -20°C until use. 50 µM guide RNA (gRNA) was prepared by mixing equal volumes of 100 µM crRNA and 100 µM tracrRNA, followed by annealing for 5 min at 95°C, cooling down at 0.1°C/sec to 25°C, incubating for 5 min at 25°C, and rapid cooling down to 4°C (stored at -20°C until use). Injection mixes were prepared fresh on the day of micro-injections by mixing 1 µL *kita* gRNA, 1 µL *rreb1a* gRNA and 1 µL *rreb1b* gRNA with 2.4 µL Alt-R® Cas9 (IDT) and 4.6 µL ultra-pure $H_2O$ for crispants; or 1 µL *kita* gRNA, 0.8 µL Alt-R® Cas9 and 7.2 µL ultra-pure $H_2O$ for controls. Mixes were incubated at 37°C for 5 min and 1 µL of phenol red (Sigma-Aldrich, MERCK, Sweden) was added as a visual injection aid. For micro-injections, eggs from all clutches of the same round of crossings of several adults were mixed and then split into two groups to generate *rreb1a*/*b* crispants and controls. Micro-injections into the cell or into the yolk close to the cell were performed at the single-cell stage using standard microinjection equipment.

At 1 dpf, dead embryos and unfertilized eggs were removed and the remaining embryos were aliquoted at 50 to 60 embryos per petri dish. Eggs and embryos were kept in filtered water with methylene blue until 5 dpf. At 4 or 5 dpf, the success rate of CRISPR/Cas9 was assessed by optically checking for lack of pigmentation using a stereo microscope, since loss of *kita* results in absence and/or reduced migration of melanocytes [54, 55]. Larvae with pigmentation were discarded. Across the individual experiments performed to reach the final sample size, 116 to 424 larvae per experiment survived to day 5. Survival from day 1 to 5 was on average 41 ±18%

(mean±SD) in *rreb1a/b* crispants and 64±13% in controls. While this reflects a significant difference in survival rate between *rreb1a/b* crispants and controls (P = 2E-3), the difference is not dissimilar to what we observed across experiments for 67 candidate genes for cardiometabolic diseases targeted the same way, with average survival rates of 37% in crispants for candidate genes and 58% in controls (unpublished data). Thus, mutations in *rreb1a* and *rreb1b* are not more harmful for early-stage development in zebrafish than mutations in other candidate genes for cardiometabolic diseases.

At 5 dpf, non-melanized embryos from each group were placed in experimental tanks at a ratio of 70% *rreb1a/b* crispants and 30% controls, at 30 or 60 embryos per tank, in 300 or 600 mL of filtered water, respectively. Larvae were fed twice daily at approximately 9AM and 3:30PM, on a standardized amount (16 mg/30 larvae) of regular dry food per feeding (zebrafeed < 100 μm, SPAROS Lda, Portugal). Feeding started in the afternoon of day 5 and was continued until the afternoon of day 9. Full water exchanges were performed midday at 7 and 9 dpf. The experiment was performed a total of six times.

While we were able to target all four putative transcripts of the zebrafish *rreb1a* with a single sgRNA, this was not possible for *rreb1b*. In *rreb1b*, we were able to target two major transcripts that code for 1499 and 1671 amino acid long proteins, while the putative transcript rreb1b-202 could not be targeted. However, this transcript has only the 5'UTR in common with one of the other transcripts and its coding region encodes a peptide that is only 29 amino acids long and does not align to any of the other amino acid sequences. Therefore, we conclude that rreb1b-202 does not code for a functional *rreb1b* isoform and, therefore, does not need to be targeted. It may still be involved in regulatory functions but these were not the focus of this study.

*Imaging of zebrafish larvae at 10 dpf*

In the morning of 10 dpf, live zebrafish larvae were washed twice with filtered water, followed by incubation for 30 min at 28.5°C in 12.5 µM monodansylpentane (MDH, SM1000a; ABCEPTA, USA) [56] in PBS (0.8 mL PBS per larva) to label neutral lipids. Next, larvae were anesthetized by adding tricaine to a final concentration of 230 µM, and were then placed in a 96-well plate. From here they were automatically aspirated, positioned and oriented in a glass capillary using an Autosampler and Vertebrate Automated Screening Technology (VAST) BioImager (Union Biometrica, Belgium) that is built on the stage of a Leica DM6000B fluorescence microscope with a Leica DFC 365 FX CCD camera (Micromedic AB, Sweden). In each larva, 12 full body images were first acquired; one every 30° of rotation around the longitudinal axis of the body using the VAST bioImager's bright field camera. Optical sections of the pancreatic islet (TexasRed filter set, HCX APO LU-V-I 40x/0.80 WATER objective, 45 images/stack, 66.04 µm Z-size) and liver (GFP and CFP filter sets, HC APO LU-V-I 10x/0.3 WATER objective, 35 images/stack, 51.03 µm Z-size) were then acquired using the fluorescence microscope. After imaging, larvae were dispensed back into 96-well plates, sacrificed by prolonged exposure to tricaine, and kept on ice. Water was removed and samples were stored at -20°C until further analysis.

*Fragment length analysis and qPCR in zebrafish larvae*

PCR products were diluted 1:10 to 1:20 before mixing 1.5 µL of diluted sample with 10 µL Hi-Di buffer (containing 73.3x diluted GS400HD ROX standard (ThermoFisher Scientific, Sweden)), followed by denaturation at 95°C for 5 min, rapid cool down on ice, and capillary electrophoresis on a DNAnalyzer (3730xl, ThermoFisher Scientific, Sweden). Chromatograms were analyzed using Peak Scanner™ software v1.0 and v2.0 (ThermoFisher Scientific, Sweden) and fragment lengths were exported for further analysis with a custom markdown script in R that calculates the relative area of the peak at the wild type allele´s fragment length relative to areas of any other peaks within ±50 base pairs of the wild type peak. PCR products generated from DNA of un-injected larvae from the same crossing were used to experimentally determine the fragment length and relative peak area of the *rreb1a* and *rreb1b* wild type PCR product.

To ascertain how well fragment length analysis managed to quantify CRISPR/Cas9-induced mutagenesis, we additionally used DNA from a subset of experimental larvae (n=126) and un-injected controls (n=32) for a qPCR-based analysis [57]. qPCR was performed in duplicate per sample, in 10 µL reactions using 1-2 µL of 10-20x diluted DNA as a template and 200-400 nM primers (**ESM Table 3**) with the PowerUP™ SYBR™ Green Master mix (ThermoFisher Scientific, Sweden) in a AriaMx Real-Time PCR System (Agilent Technologies, USA). Dilution series of samples from un-injected siblings were used as a reference for relative quantification. Two non-targeted loci in *rreb1a* and *rreb1b* were used for relative quantification of genomic DNA and normalization of quantification data from the on-target primer pairs. Based on the congruence across qPCR and fragment length analysis results in un-injected and injected larvae, samples were assigned to the control group if >70% of the *rreb1a* peak area and >60% of the *rreb1b* peak area was wildtype; while larvae with ≤60% and ≤50% of peak areas being wildtype for *rreb1a* and *rreb1b* were assigned to the *rreb1a/b* crispant group (**ESM Fig. 7**). Larvae that did not fulfill either criterion were excluded from the analysis. Across all six rounds of the experiment, a total of 175 larvae were imaged at 10 dpf, and 49 additional 10 dpf larvae were characterised using biochemistry only, due to time constraints on the day of imaging. Of these 224 larvae, 92 were *rreb1a/b* crispants; 64 were controls; and 68 had inconclusive fragment length analysis results and were excluded from the analysis.

*Zebrafish statistical analysis*

While all outcomes were normally distributed, inverse normal transformations were applied before the statistical analysis to enable a comparison of effect sizes across outcomes. Genetic effects on outcomes of interest were subsequently examined using linear regression analysis, adjusting for the time of day at which larvae were imaged; the tank larvae were in from day 5 to 10; and the round of the experiment in which they were examined (1-6). Effects on dorsal and lateral body surface area were additionally adjusted for body length as a covariable; and effects on whole-body glucose, LDLc, triglyceride and total cholesterol levels were additionally adjusted for the position of the sample in the Mindray Analyser; the run in which the sample was analyzed; and whether the larva had been imaged or not.

*Generation of RREB1$^{KO/KO}$ EndoC-βH1 cells*

sgRNAs were taken from the TKO Library v3 [58] or designed with the CRISPOR online design tool (http://crispor.tefor.net) [59]. sgRNA oligonucleotides targeting *RREB1* exon 4 (ATGACGTCAAGTTCGCCCGC), exon 5 (AGTGCAAATCTTCTCACACA), exon 8 (GTATGGACTGGAGACCCACA), and exon 12 (GACAGACTCCCCCAAAAGCG) were amplified [58], and sub-cloned into the *BsmBI* restriction enzyme sites in the lentiviral vector plentiCRISPRv2 [60]. pLentiCRISPRv2 was a gift from Feng Zhang (http://n2t.net/addgene:52961; RRID: Addgene_52961). Lentiviruses were produced by Lenti-X HEK293T cells co-transfected with 6.85 μg of pMD2.G (RRID:Addgene_12259), 10.3 μg of psPAX2 (RRID:Addgene_12260) and 12.85 μg pLentiCRISPRv2-sgRNAs. Functional viral titer was performed in EndoC-βH1 cells as described previously [26].

*EndoC-βH1 insulin secretion assays*

Live cell count was quantified using the fluorescent based CyQUANT Direct Cell Proliferation Assay (ThermoFisher Scientific, UK). Cellular insulin content was extracted using ice-cold acid-ethanol (1.5% concentrated HCl, 75% ethanol and 23.5% deionized water). The Insulin (human) AlphaLISA Detection Kit (PerkinElmer, USA/UK) was used to measure the amount of secreted insulin (supernatants) and cellular insulin content. Samples were diluted in 1x AlphaLISA immunoassay buffer (supernatant 1:10, content 1:50) and analysis was performed in white 96-well 1/2 Area Plates (PerkinElmer, USA/UK) on the EnSpire plate reader. Sample values were interpolated from an insulin analyte standard curve included on every plate using a four-parameter non-linear regression of log-transformed insulin count data in Prism 8 (GraphPad Software).

*Genome editing of hiPSCs*

*BsbI* restriction enzyme-mediated sub-cloning of annealed gRNA oligonucleotides into plasmid pX330-U6-Chimeric_BB-CBh-hSpCas9 [61] as previously described [62]. pX330-U6-Chimeric_BB-CBh-hSpCas9 was a gift from Feng Zhang (RRID:Addgene_42230). Human iPSCs were either co-transfected with 1 μg of pX330-Puro-Cas9 plasmid containing exon 10 sgRNA and 100 nM of the repair template (*RREB1*$^{WT/WT}$) or with pX330-Puro-RREB1 plasmids containing exon 4 and exon 10 sgRNAs (*RREB1*$^{KO/KO}$) using FuGENE® 6 (Promega, UK) before puromycin treatment (400 ng/mL) to select for transfected cells. Selection media was removed after 48 hours and hiPSCs were grown in antibiotic-free mTeSR1 until ~90% confluency. Cells were then re-plated at low density (2,000 cells/60 mm dish), allowed to form clones, and were picked using a microscope-mediated pipetting approach into individual wells of a 96-well plate containing mTeSR1. After approximately seven days, colonies were split into two replica pre-coated 96-well plates for either expansion or genotyping.

*Differential expression and functional enrichment analysis*

Reads were mapped to the human genome build hg19 using STAR v.2.5 [63]. GENCODE v19 (https://www.gencodegenes.org/releases/19.html) was used as a transcriptomic reference [64]. Quantification of gene expression was performed by featureCounts from the Subread package v.1.5 (http://subread.sourceforge.net/) [65]. To adjust for technical effects, removal of unwanted variation was conducted in R using the RUVSeq following the instructions in the manual compiled on May 2, 2019 [66]. Read counts were filtered to include only genes that reached one transcript per million (TPM) in at least one cell line and in at least one stage before normalisation. Differential gene expression analysis was conducted in R using the Bioconductor package DESeq2 [67] according to the vignette compiled on November 30, 2016. Differentially expressed genes were evaluated for enrichment in gene ontology terms (GO biological processes, KEGG and Reactome pathways) using g:Profiler with the significance threshold set to $p<0.01$ using the tailor-made g:SCS algorithm for multiple testing (https://biit.cs.ut.ee/gprofiler/gost) [68]. Correlation of gene expression patterns during beta cell development was calculated following the weighted gene co-expression network analysis (WGCNA) R software package (v.1.51) [38, 39]. Differential *RREB1* exon usage was analysed using the Bioconductor package DEX-Seq [69].

### ESM Tables

#### ESM Table 1: *RREB1* orthologues in *Danio rerio* (Ensembl release 105 - Dec 2021)

| | Orthologue 1 | Orthologue 2 |
|---|---|---|
| **Gene** | *rreb1a* | *rreb1b* |
| **Ensembl gene ID** | ENSDARG00000063701 | ENSDARG00000042652 |
| **Type** | 1-to-many | 1-to-many |
| **% identity of zebrafish sequence to human sequence** | 45.30% | 42.73% |
| **% identity of human sequence to zebrafish sequence** | 44.49% | 40.99% |
| **Gene Order Conservation (GOC) Score; 0-100** | 50 | 25 |
| **Whole Genome Alignment Coverage (WGA); 0-100** | 74.18 | 73.19 |
| **High Confidence** | yes | yes |

**ESM Table 2: Gene IDs, CRISPOR scores and gRNA target sequences for zebrafish genes *rreb1a*, *rreb1b* and *kita*.**

| **ENSEMBL gene ID** | **gene name** | **CRISPOR scores** [azimuth in-vitro; off-targets with 0-1-2-3-4 mismatches; (off-targets in 12 nts next to PAM)] | **target sequence (DNA)** |
|---|---|---|---|
| ENSDARG00000063701 | rreb1a | 53; 0-0-1-3-65; (0-0-0-0-0) | CGTTGGTGGTGAAGGTCTGA |
| ENSDARG00000042652 | rreb1b | 52; 0-0-0-5-43; (0-0-0-0-3) | GGTCAGACATTCACCACCAA |
| ENSDARG00000043317 | kita | 74; 0-0-0-2-8; (0-0-0-0-0) | GGCTGACCCAGTGTGATCGG |

### ESM Table 3: Sequences of primers used for fragment length analysis

| F-primer name | seq 5´to 3´ | product size |
|---|---|---|
| rreb1a_33-T1 F | tgtaaaacgacggccagtCCAAGTTGGTTTTTCCTCCA | 345 |
| rreb1a_33-T1 R | gtgtcttTGGCAGGGCAAGTAAAAATC | |
| rreb1b_33-T1 F | tgtaaaacgacggccagtAGCTCTCTGGACCGACACAT | 204 |
| rreb1b_33-T1 R | gtgtcttATAAAAAGCCCATGCAGCTC | |

Lower case characters in F primers are M13-F primer binding sites and PIG-tail sequences in R primers

**ESM Table 4: Sequences of primers used for qPCR based analysis of mutagenesis efficiency.**

| F-primer name | seq 5´to 3´ (“/” indicates the predicted cut site in a target) | product size |
|---|---|---|
| rreb1a-qPCRgD_T33-F2 | CGTGCAGTATCTGCGGGAAG | 121 bp |
| rreb1a-qPCRgD_T33-R2 | CCGTTGGTGGTGAAGGT/CTG | |
| rreb1a-qPCRgD_Ex11-F2 | CCTATGGGTGAGGCAATGGA | 109 bp |
| rreb1a-qPCRgD_Ex11-R2 | GCTCTACCACGGCTGCCTCT | |
| rreb1b-qPCRgD_T33-F1 | CGGTCAGACATTCACCA/CCA | 112 bp |
| rreb1b-qPCRgD_T33-R1 | AAGCCCATGCAGCTCAACAA | |
| rreb1b-qPCRgD_Ex10-F1 | TCCTAGCCTTCCACCCCAAA | 98 bp |
| rreb1b-qPCRgD_Ex10-R1 | CCCAATTGGTGGGAGAGGAG | |

T33 refers to primers at the target site; Ex11 and Ex10 refer to primers outside the target site.

**ESM Table 5: Ten most common coding mutations in TP53 that were assessed in genome-edited hiPSC lines.**

| Common *TP53* mutations |
|---|
| P151S |
| R175H |
| R181H |
| H193R |
| M237I |
| G245S |
| R248W |
| R267W |
| R273L |
| R273G |

### ESM Table 6: Composition of basal and complete differentiation media.

| Basal differentiation medium | | | | |
|---|---|---|---|---|
| **Stage 1 and 2** | **Stage 3 and 4** | **Stages 5-7** | **Manufacturer** | **Catalog #** |
| MCDB 131 | MCDB 131 | MCDB 131 | ThermoFisher Scientific | 10372019 |
| 0.1% P/S | 0.1% P/S | 0.1% P/S | Sigma-Aldrich | P0781 |
| 1.5 g/L $NaHCO_3$ | 2.5 g/L $NaHCO_3$ | 1.5 g/L $NaHCO_3$ | ThermoFisher Scientific | 25080060 |
| 1x Glutamax | 1x Glutamax | 1x Glutamax | ThermoFisher Scientific | 35050038 |
| 10 mM Glucose | 10 mM Glucose | 20 mM Glucose | ThermoFisher Scientific | A2494001 |
| 0.5% BSA | 2% BSA | 2% BSA | Roche | 10775835001 |
| | 1:200 ITS-X* | 1:200 ITS-X* | ThermoFisher Scientific | 51500056 |
| | | 10 µM Zinc sulfate | Sigma-Aldrich | Z0251 |

*Insulin-Transferrin-Selenium-Ethanolamine
**3,3,5-Triiodo-L-thyronine sodium salt

**Complete differentiation medium**

| Stage | Medium | Day | Factors | Concentration | Manufacturer | Cat # |
|---|---|---|---|---|---|---|
| Definitive endoderm (DE) | MCDB 131 (Stage 1 and 2) | 1 | CHIR 99021 | 3 µM | Axon Medchem | 1368 |
| | | | Activin A | 100 ng/mL | PeproTech | 120-14 |
| | | 2 | CHIR 99021 | 0.3 µM | Axon Medchem | 1368 |
| | | | Activin A | 100 ng/mL | PeproTech | 120-14 |
| | | 3 | Activin A | 100 ng/mL | PeproTech | 120-14 |
| Primitive gut tube (PGT) | MCDB 131 (Stage 1 and 2 | 4-5 | KGF | 50 ng/mL | PeproTech | 100-19 |
| | | | Ascorbic acid | 0.25 mM | Sigma-Aldrich | A4544 |
| Posterior foregut (PFG) | MCDB 131 (Stage 3 and 4) | 6-7 | Retinoic acid | 1 µM | Sigma-Aldrich | R2625 |
| | | | Sant-1 | 0.25 µM | Sigma-Aldrich | S4572 |
| | | | KGF | 50 ng/mL | PeproTech | 100-19 |
| | | | LDN193189 | 100 nM | Stemgent | 04-0074 |
| | | | PKC act V (TBP) | 200 nM | Merck | 565740 |
| | | | Ascorbic acid | 0.25 mM | Sigma-Aldrich | A4544 |
| Pancreatic endoderm (PE) | MCDB 131 (Stage 3 and 4) | 8-10 | Retinoic acid | 0.1 µM | Sigma-Aldrich | R2625 |
| | | | Sant-1 | 0.25 µM | Sigma-Aldrich | S4572 |
| | | | KGF | 2 ng/mL | Peprotech | 100-19 |
| | | | LDN193189 | 200 nM | Stemgent | 04-0074 |
| | | | PKC act V (TBP) | 100 nM | Merck | 565740 |
| | | | Ascorbic acid | 0.25 mM | Sigma-Aldrich | A4544 |
| Endocrine progenitors (EP) | MCDB 131 (Stage 5-7) | 11-13 | Retinoic acid | 0.05 µM | Sigma-Aldrich | R2625 |
| | | | Sant-1 | 0.25 µM | Sigma-Aldrich | S4572 |
| | | | LDN193189 | 100 nM | Stemgent | 04-0074 |
| | | | ALK5 Inhibitor II | 10 µM | Enzo Life Sciences | ALX-270-445 |
| | | | T3** | 1 µM | Sigma-Aldrich | T6397 |
| | | | Heparin sodium salt | 10 µg/mL | Sigma-Aldrich | H3149 |
| Endocrine cells (EN) | MCDB 131 (Stage 5-7) | 14-20 | LDN | 100 nM | Stemgent | 04-0074 |
| | | | ALK5 Inhibitor II | 10 µM | Enzo Life Sciences | ALX-270-445 |
| | | | T3** | 1 µM | Sigma-Aldrich | T6397 |
| | | | Heparin sodium salt | 10 µg/mL | Sigma-Aldrich | H3149 |
| | | | y-Secretase Inhibitor XX | 100 nM | Merck | 565789 |
| Beta-like cells (BLC) | MCDB 131 (Stage 5-7) | 21-27 | ALK5 Inhibitor II | 10 µM | Enzo Life Sciences | ALX-270-445 |
| | | | T3** | 1 µM | Sigma-Aldrich | T6397 |
| | | | N-acetyl-Cys | 1 mM | Sigma-Aldrich | A9165 |
| | | | Trolox | 10 µM | Merck | 648471 |
| | | | R428 | 2 µM | Selleck Chemicals | S2841 |
| | | | Heparin sodium salt | 10 µg/ml | Sigma-Aldrich | H3149 |

### ESM Figures

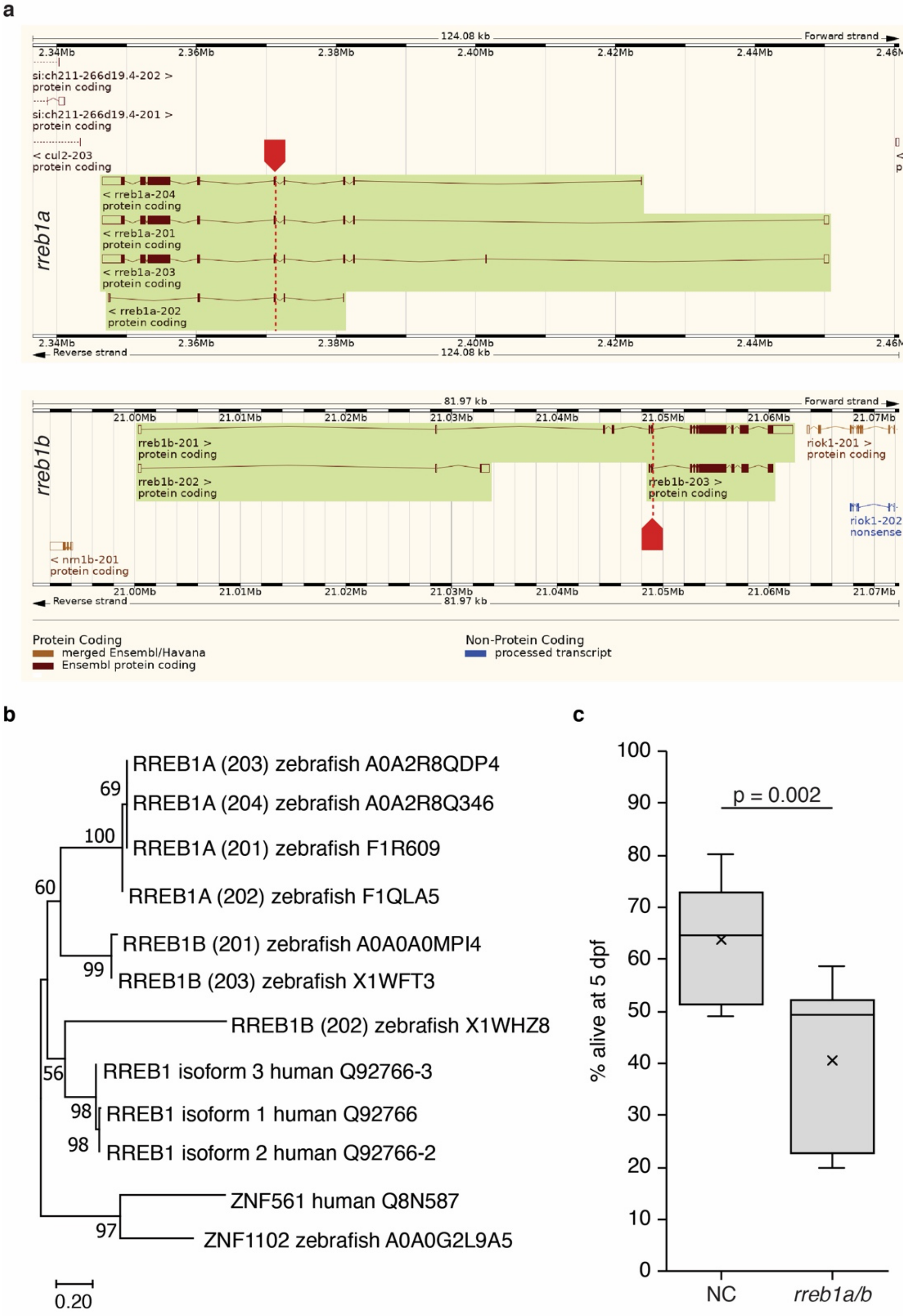


**ESM Fig. 1: Characterisation of zebrafish model.**
(**a**) Genomic structure of the zebrafish orthologues (*rreb1a* and *rreb1b*) of human *RREB1* and the sites targeted by CRISPR/Cas9 in each gene. (**b**) Phylogenetic tree of human and zebrafish RREB1 proteins. Numbers on nodes are bootstrap values and uniprot accession numbers of the sequences used are given after the species name. (**c**) Box and whisker plot of the percentage of embryonic/larval survival from day 1 to 5 post-fertilization for the control group (NC) and *rreb1a/b* crispants based on data from six independent experiments (the number of larvae per experiment ranged from 188 to 308 control and mutant larvae 24 h after micro-injections; 1806 larvae in total). (Paired Student’s t-test)

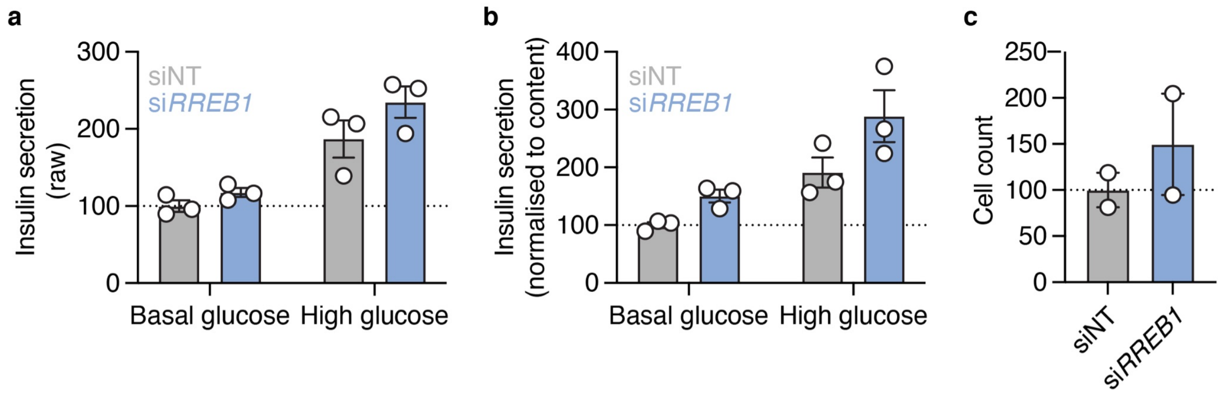


**ESM Fig. 2: Characterization of glucose-stimulated insulin secretion from si*RREB1* EndoC-βH1 cells.**

(**a**) Raw glucose stimulated insulin secretion (% of *RREB1*$^{WT/WT}$ basal glucose) for siNT and si*RREB1* cells. (**b**) Glucose stimulated insulin secretion normalised to insulin content (% of siNT basal glucose) for siNT and si*RREB1* cells. (**c**) Cell count measurements used to normalize glucose stimulated insulin secretion. (Unpaired t-test)

**a**

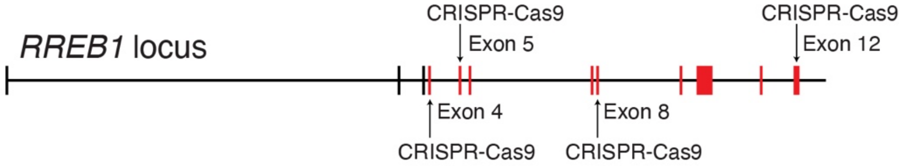


**b**

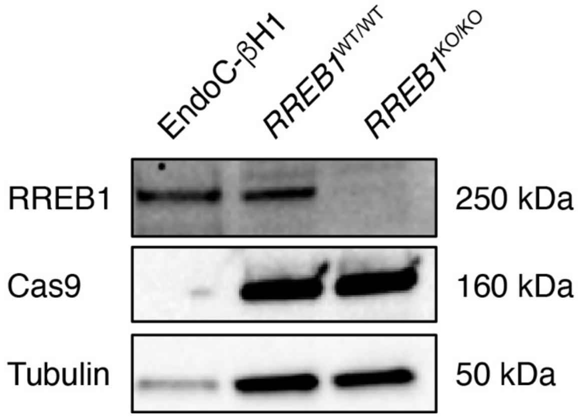


**ESM Fig. 3: Generation and characterisation of *RREB1*$^{KO/KO}$ EndoC-βH1 cells.** (**a**) Schematic highlighting the genomic locations of the four sgRNAs used to generate *RREB1*$^{KO/KO}$ EndoC- βH1 cells. (**b**) Western blot for RREB1 (250 kDa), Cas9 (160 kDa), and Tubulin (50 kDa) in parental EndoC-βH1, *RREB1*$^{WT/WT}$ control cells, and *RREB1*$^{KO/KO}$ knockout cells.

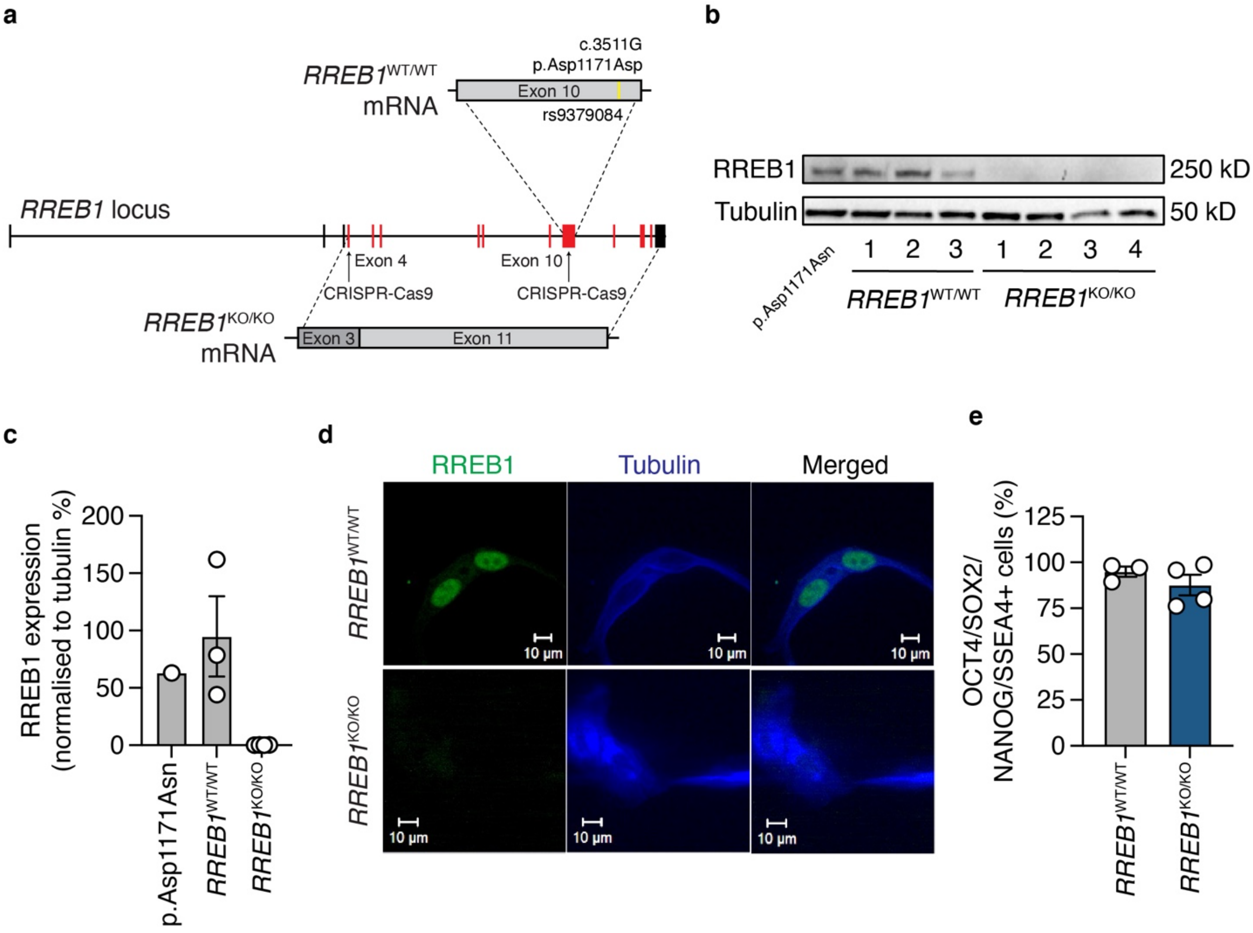


**ESM Fig. 4: Generation and characterisation of *RREB1*$^{KO/KO}$ hiPSC lines.**
(**a**) Schematic highlighting the genomic locations of the two sgRNAs used to generate *RREB1*$^{KO/KO}$ hiPSC lines by deleting ~50 kb of the *RREB1* gene. Located in exon 10 is rs9379084 (yellow) that was genetically engineered to be homozygous for the major p.Asp1171 allele. (**b-c**) Western blot and quantification for RREB1 (250 kDa) and Tubulin (50 kDa) in p.Asp1171Asn, *RREB1*$^{WT/WT}$ and *RREB1*$^{KO/KO}$ hiPSC lines. (**d**) Immunofluorescence staining of RREB1 (green) and tubulin (blue) in *RREB1*$^{WT/WT}$ and *RREB1*$^{KO/KO}$ hiPSC lines. (**e**) Quantification of percent of hiPSC cells expressing pluripotency proteins OCT4, SOX2, NANOG and SSEA4.

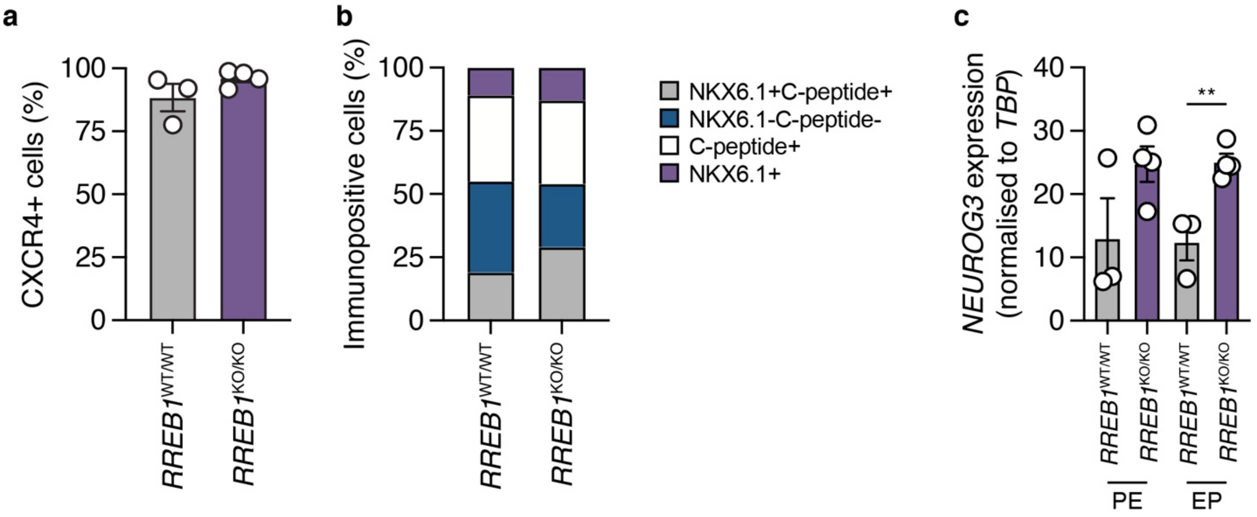


**ESM Fig. 5: Characterisation of *in vitro* differentiation of *RREB1*$^{KO/KO}$ and *RREB1*$^{WT/WT}$ clones towards beta-like cells.**

(**a**) Flow cytometric quantification of CXCR4+ definitive endoderm cells derived from *RREB1*$^{KO/KO}$ and *RREB1*$^{WT/WT}$ hiPSC lines. (**b**) Proportion BLCs of derived from *RREB1*$^{WT/WT}$ and *RREB1*$^{KO/KO}$ hiPSC cells immunopositive for NKX6.1+C-peptide+, NKX6.1-C-peptide-, C-peptide+, or NKX6.1+. (**c**) Expression of *NEUROG3* transcript in *RREB1*$^{KO/KO}$ and *RREB1*$^{WT/WT}$ hiPSC-derived pancreatic endoderm (PE) and endocrine progenitor (EP) cells. n=3-4. Data are presented as mean±SEM. p**<0.01. (Unpaired t-test)

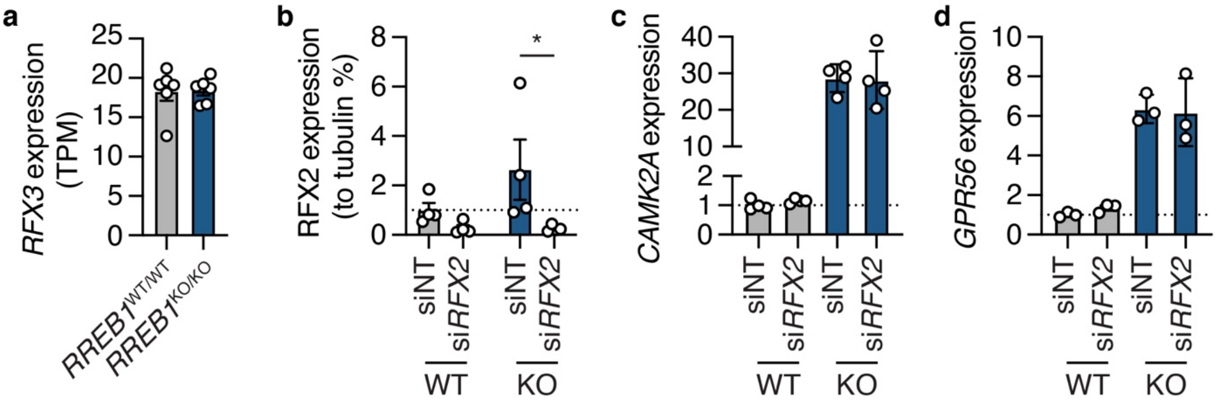


**ESM Fig. 6: RREB1 deficiency in EndoC-βH1 cells alters gene expression of *RFX* family members.**

(**a**) Gene expression of *RFX3* in *RREB1*$^{WT/WT}$ and *RREB1*$^{KO/KO}$ EndoC-βH1 cells. (**b**) RFX2 protein quantification normalised to tubulin in *RREB1*$^{WT/WT}$ and *RREB1*$^{KO/KO}$ EndoC-βH1 cells following siNT and si*RFX2* transfection. (**c-d**) Gene expression of (**c**) *CAMK2A* and (**d**) *GPR56* (% of siNT, normalised to housekeeping gene) in *RREB1*$^{WT/WT}$ and *RREB1*$^{KO/KO}$ EndoC-βH1 cells following siRNA-mediated deletion of *RFX2*. n>3. Data are presented as mean±SEM. *p<0.05. (Unpaired t-test)

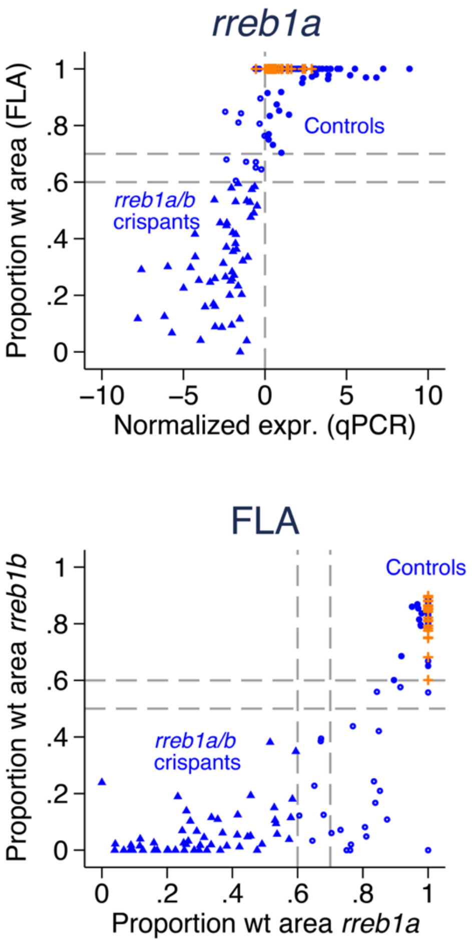


**ESM Fig. 7: Results from fragment length analysis and qPCR for 126 microinjected zebrafish larvae (10 dpf) and 32 un-injected controls.**
Comparing results from fragment length analysis and qPCR performed on the same samples shows that un-injected controls (orange crosses) have 100% of rreb1a (top left) and >60% of rreb1b (top right) peak area at the expected wildtype length. For both zebrafish genes, normalised gene expression based on qPCR results (i.e. residuals of expression at the CRISPR-targeted site adjusted for expression at a later exon) is >0 for most samples. Based on these results, we used fragment length analysis results for both genes (bottom left) to allocate larvae to the control group (blue triangles) if >70% of the rreb1a peak area and >60% of the rreb1b peak area was wildtype; while larvae with ≤60% and ≤50% of peak areas being wildtype for rreb1a and rreb1b were assigned to the rreb1a/b crispant group (filled blue circles). Larvae not fulfilling either criterion were excluded from the analysis (open blue circles).

### ESM References
